# Accurate identification of structural variations from cancer samples

**DOI:** 10.1101/2023.05.31.543104

**Authors:** Le Li, Chenyang Hong, Jie Xu, Claire Yik-Lok Chung, Alden King-Yung Leung, Delbert Almerick T. Boncan, Lixin Cheng, Kwok-Wai Lo, Paul B. S. Lai, John Wong, Jingying Zhou, Alfred Sze-Lok Cheng, Ting-Fung Chan, Feng Yue, Kevin Y. Yip

## Abstract

Structural variations (SVs) are commonly found in cancer genomes. They can cause gene amplification, deletion, and fusion, among other functional consequences. With an average read length of hundreds of kilobases, nano-channel-based optical DNA mapping is powerful in detecting large SVs. However, existing SV calling methods are not tailored for cancer samples, which have special properties such as mixed cell types and sub-clones. Here we propose the COMSV method that is specifically designed for cancer samples. It shows high sensitivity and specificity in benchmark comparisons. Applying to cancer cell lines and patient samples, COMSV identifies hundreds of novel SVs per sample.

## 1 Background

Structural variations (SVs), such as large insertions, deletions, inversions, translocations, and copy number variations, are commonly found in cancer genomes [1–3]. Their high prevalence is due to a combination of factors, including defects in the DNA replication and repair pathways and inefficient apoptotic response [4–7]. SVs in cancer genomes can cause a variety of functional consequences, including altered protein sequence, gene loss, change of gene copy number, gene fusion, perturbed epigenetic signals and gene regulation, and change of 3D genome structure [8–11]. Furthermore, some SVs are recurrent in particular cancer types [12–14], rendering them potentially useful for cancer diagnosis and classification.

It is challenging to detect large SVs by short read sequencing due to difficulties in read alignment and determination of the whole genomic span affected by an SV, especially when the break points are within tandem repeats or when the SV involves DNA contents not contained in the reference [15, 16]. As such, it has been shown advantageous to use different experimental technologies to detect SVs of different sizes and complexities [10]. Among the technologies currently available, optical DNA mapping [17, 18], due to its long read length (hundreds of kilobases on average), is powerful for detecting large SVs. The low cost also makes optical DNA mapping an attractive option as compared to long-read sequencing.

In nano-channel-based optical mapping systems [18], specific sites on the DNA are labeled either by single-stranded enzymatic nicking followed by repair with fluorescent dye conjugated nucleotides, or by direct labeling methods without nicking DNA (e.g, Bionano Genomics’s Direct Label and Stain [DLS] protocol). The labeled DNA molecules are linearized and imaged in nanometer-scaled channels. The positions of the fluorescent labels on each molecule define a signature of the molecule, which can be compared with the expected label positions derived from a reference genome to determine the genomic location from which the molecule originated and identify any differences between the DNA molecule and the aligned region of the reference.

Based on this idea, a few methods have previously been proposed for calling SVs from optical mapping data [19–21]. These methods have been extensively applied to identify SVs from human samples [22–24]. However, all these methods were originally developed for non-cancer samples, and are thus not suitable for cancer samples due to their unique properties, as follows.

First, cancer samples usually contain a mixture of other cell types in addition to cancer cells, such as cells from adjacent non-cancer tissues, immune cells, and stromal cells [25]. As a result, for an SV specific to cancer cells, among all the DNA molecules from that locus, only those coming from the cancer cells would contain information about the SV. This fraction of SV-supporting molecules can be small when the tumor content in the sample is low, and it is further halved for a heterozygous SV. An SV calling method developed for non-cancer samples can easily miss these SVs if it wrongly assumes that all the cells have the same genetic makeup and expects each SV to have around 100% or 50% of the aligned molecules supporting it in the cases of a homozygous and heterozygous SV, respectively. This issue is further aggravated by the presence of cancer sub-clones, each of which can contain a different set of SVs.

Second, it is common for cancer genomes to have abnormal copy numbers at different scales, from specific focal genomic regions (e.g., extrachromosomal circular DNAs) to whole chromosomes (aneuploidy). Therefore, the proportion of SV-supporting molecules can vary substantially along the genome. An SV calling method designed for non-cancer samples would fail to detect many SVs if it wrongly assumes that the proportion of SV-supporting molecules distributes around two constant mean values, *x*% and *x/*2% respectively for all homozygous and heterozygous SVs in the whole genome, even if it allows *x* to take a value other than 100.

Third, as compared to non-cancer genomes, cancer genomes usually contain substantially more SVs and the SVs are more complex. Accordingly, the resulting alignments and *de novo* assemblies of optical mapping data usually contain more errors, which can lead to inaccurate SV calls for SV identification methods that depend heavily on the reliability of the alignments or assemblies.

Realizing these challenging properties of cancer genomes, here we propose a new method, COMSV (Cancer Optical Mapping for detecting Structural Variations), which is specifically designed for cancer samples. We show that it has higher precision and sensitivity as compared to existing SV callers. We also demonstrate the use of COMSV in accurately calling SVs from both cancer cell lines and patient tumor samples.

In addition, we propose a new scheme for SV annotation that provides more information than the ones used by the previous SV calling methods, taking into account redundancy of reported SVs (e.g., not to report a duplication also as an insertion), uncertainty of SV break points (e.g., distinguishing between fully identified SVs and those having only partial break point information), and potential functional consequences of the SVs.

## 2 Results

### 2.1 The COMSV method

COMSV includes a pipeline for identifying insertions and deletions (the “indel pipeline”) and a pipeline for identifying other types of SVs (the “complex SV pipeline”). A high-level description of the pipelines is given below, whereas the details are provided in Methods.

The indel pipeline, which is based on alignment results, consists of four main steps (Figure 1a-d), namely 1) processing of the resulting alignments, 2) extraction of inter-label distances, 3) clustering of the inter-label distances and initial identification of indels, and 4) post-processing of the indel calls. Each of these steps involves designs customized for cancer samples.

**Figure 1:**
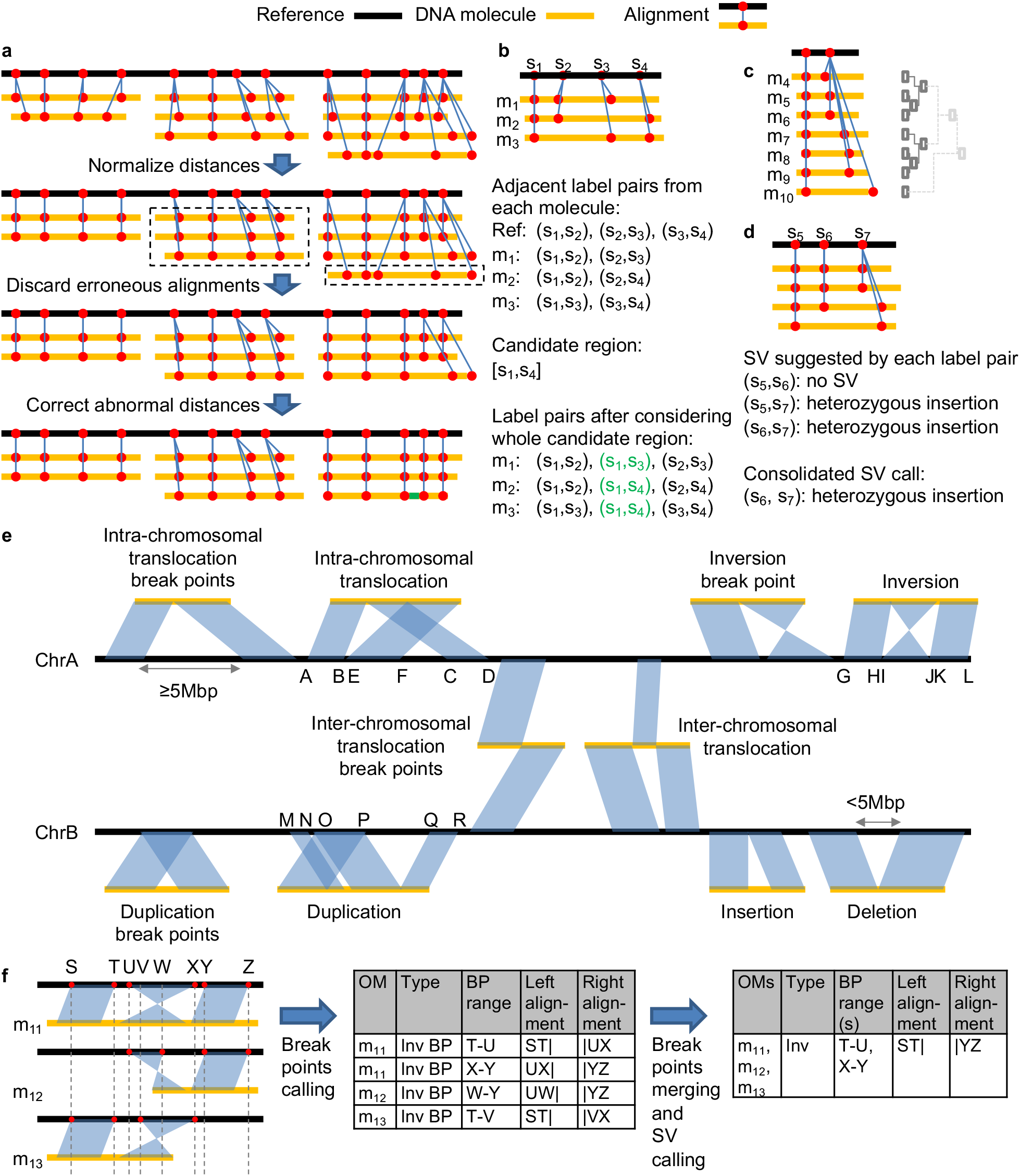
The COMSV pipelines, which include the indel pipeline (a-d) and the complex SV pipeline (e-f). **a** Observed-to-expected distance ratios between two labels can deviate from 1 due to scaling or alignment errors (first row). After normalization, for alignments that still have these abnormal ratios (second row, dashed rectangles), the mutually reinforcing ones are considered indel candidates while the others are discarded (third row). Isolated label pairs with abnormal distances are instead corrected (fourth row, green bar). **b** Adjacent label pairs are collected from individual molecules to form candidate indel regions based on their overlaps, which enables the collection of distances between non-adjacent labels from a molecule if they are adjacent in another molecule (shown in green). **c** For each label pair, the collected distances are clustered to identify the number of distinct alleles and the number of molecules that support each of them. **d** For each potential indel region, the clustering result of each label pair suggests an initial SV type, and the final SV type of the whole region is determined by considering these suggestions jointly. **e** Different types of SVs that can be identified from split-alignments. An SV break point is called if a split alignment suggests an SV event, while a complete SV is called if the full span of the SV can be determined. **f** An example of calling SV break points and complete SVs. The reference is repeatedly displayed multiple times to show the split-alignment of each molecule clearly. Vertical dotted lines mark key aligned labels that define the boundary of each aligned segment on the reference. The corresponding labels on the reference are shown in red. Break point information is first collected from individual molecules and then considered together to refine the break point locations and determine whether complete SVs can be called.

COMSV takes as input the alignment of optical mapping data of individual DNA molecules to a reference map produced by the *in silico* digestion of a reference genome. In the first step (Figure 1a), COMSV normalizes the distances between labels on the molecules to correct for incomplete molecular linearization [21]. After normalization, if the distance between two adjacent labels on a molecule deviates substantially from that on the reference, there could be an indel or the alignment could be wrong. Depending on whether multiple molecules display similar distance deviations consistently, COMSV either identifies the region as having a potential indel, considers the alignment as wrong and discards it, or corrects the specific distance value according to the consensus. These three alternative treatments are crucial for cancer data, which have a relatively high error rate in the optical map alignments.

In the second step (Figure 1b), all pairs of adjacent labels on the reference genome and on the aligned molecules are extracted. Conceptually, by comparing the observed distances between two labels on the molecules and their expected distance on the reference, insertions and deletions can be identified. However, unlike previous methods that consider each label pair separately, COMSV groups the overlapping ones into candidate indel regions and considers all pairs of labels within each region at the same time, which provides more information for accurate indel calling. Again, this design reduces the negative effects of incorrect molecule alignments to the SV calling procedure.

In the third step (Figure 1c), for each label pair, their distances on the aligned molecules are clustered. In most situations, these distances form either one single cluster or two clusters, the former of which corresponds to having no indel or a homozygous indel while the latter corresponds to having a heterozygous indel. However, due to mixed cell types, sub-clones and copy number changes, the proportion of molecules that support the indel allele is not known *a priori*, and the flexibility of the clustering approach enables highly sensitive detection of indels regardless of the exact proportions. COMSV then determines both the SV type and zygosity of each potential indel according to the clustering results.

In the fourth step (Figure 1d), the initial indels identified in the third step are further processed to remove redundancy and determine the consensus span. Each resulting indel is then given a confidence score based on a model obtained by machine learning, which helps prioritize the identified indels for experimental follow-ups.

The complex SV pipeline uses split-aligned molecules, as in previous methods, partially unaligned molecules, and contigs assembled *de novo* from individual molecules to identify SVs (Figure 1e-f). A split-aligned molecule is one with different parts of it aligned separately to different regions of the reference. Based on the locations and orientations of these different sub-alignments, the type of each candidate SV can be inferred (Figure 1e). Since it is generally difficult for alignment algorithms to make accurate split-alignments, COMSV also analyzes molecules that are partially aligned to the reference, and checks if the unaligned parts signal the possibility of an SV. For example, if the unaligned part can be aligned to the same reference location as the aligned part, there could be a tandem duplication. This re-alignment approach of COMSV can identify additional SVs missed by the split alignments. Finally, aligning contigs to the reference provides very long-range information required by some SV types such as translocations.

For all the SV candidates identified from the split-alignments and partial alignments, the information collected from different molecules is further integrated to produce a final set of non-redundant SVs with detailed annotations (Figure 1f).

We provide COMSV as an open-source tool, available freely at https://github.com/kevingroup/COMSV.

### 2.2 Comparisons between COMSV and other methods

We compared COMSV with three existing methods for identifying SVs from optical mapping data, namely Bionano Solve [19], OMIndel [20] and OMSV [21], first using simulated indels with the proportion of molecules from the SV locus that support the SV allele varying from 10% to 90% (Figure S1).

In terms of precision (i.e. the ratio of SVs called by a method that are true), all methods had stable performance when the proportion of SV-supporting molecules was reasonably large, but COMSV best maintained the high precision when the proportion of SV-supporting molecules was lower than 15%, no matter zygosity was ignored (Figure S1a) or considered (Figure S1b). This shows that the other methods were unable to distinguish between true SV cases and sizing/alignment errors when the proportion of SV supporting molecules is low.

In terms of recall (also called sensitivity, i.e., the ratio of true SVs that are called by a method), all methods performed better when the proportion of SV-supporting molecules was large, but the performance of COMSV deteriorated much more slowly than OMIndel and OMSV when the proportion decreased (Figure S1c). When correct zygosity was required, COMSV outperformed all the three other methods when the proportion of SV-supporting molecules was large (Figure S1d).

Overall, COMSV achieved the best balance between precision and recall, as indicated by its consistently high F1 score for all proportions of SV-supporting molecules (Figure S1e,f).

Next, we produced a set of simulated data with mixed cell types. Specifically, based on a cell evolution graph (Figure S2), we simulated SVs in four groups of cells, namely normal cells (Sample 1), trunk cancer cells (Sample 2), and cancer cells in two sub-clones (Samples 3 and 4). Due to the evolutionary relationships between these cells, Samples 2, 3 and 4 contain all the SVs of Sample 1, while Samples 3 and 4 contain most SVs of Sample 2. We then produced optical mapping data from each of these four pure cell types, as well as three cell mixture scenarios of increasing complexity (Table S1), namely a cancer sample with high tumor content (Sample 5), a cancer sample with low tumor content and one sub-clone (Sample 6), and a cancer sample with low tumor content and two sub-clones (Sample 7).

We evaluated the performance of the four methods based on each type of SVs they identified from each sample (Figure 2). For the homogeneous cell samples (Samples 1-4), the performance of COMSV was comparable with Bionano Solve and OMSV in calling deletions, insertions and inversions (Figure 2a-c), which is expected since Bionano Solve and OMSV were originally designed for normal samples that do not have mixed cell types. For duplications, COMSV outperformed Bionano Solve and OMSV (Figure 2d), partially attributable to its targeted re-alignment of unaligned parts of molecules.

**Figure 2:**
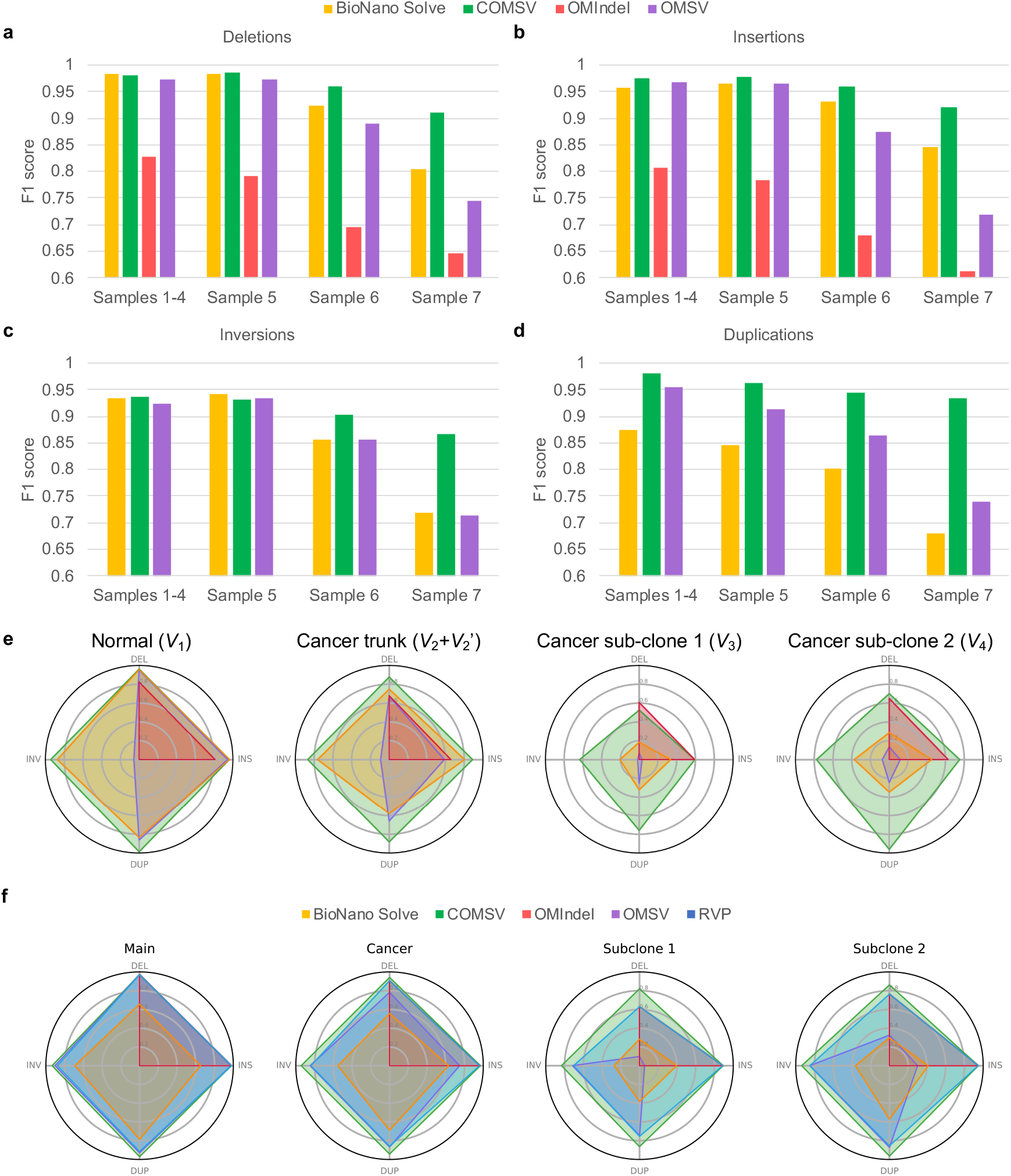
Comparisons between the performance of COMSV and three existing SV calling methods based on simulated data with different cell compositions based on a nicking enzyme (a-d) or DLS (e). **a-d** For each type of SVs, including deletions (a), insertions (b), inversions (c) and duplications (d), the performance of each method when applied to the homogeneous cell samples with one cell type per sample (Samples 1-4) and the heterogeneous cell samples with multiple cell types per sample (Samples 5-7) is shown. Since OMIndel could only identify indels, it is omitted from Panels c and d. **e-f** F1 scores of the different methods in detecting SVs in Sample 7 introduced at different stages during cancer evolution for Nt.BspQI (e) and DLS (f) labeling.

The strengths of COMSV are more clearly demonstrated by its performance on the heterogenous cell samples (Samples 5-7), especially the most complex sample with two sub-clones (Sample 7). COMSV outperformed all the other methods substantially for all four types of SVs in this sample (Figure 2a-d). When the SVs introduced at different stages during the simulated cancer evolution were investigated separately, we found that the better performance of COMSV was largely due to SVs specific to the cancer cells, especially those in the sub-clones (Figure 2e). These results show that COMSV is more capable of handling the special properties of cancer samples, such as the presence of sub-clones, than the other methods.

To investigate how the SV calling performance of the different methods was affected by alignment accuracy, we supplied the correct molecule alignments as inputs to them (Figure S3). In general, all methods benefited from receiving these correct alignments. For example, OMSV became much more capable of identifying inversions and duplications when supplied with the correct alignments (comparing Figure 2c-d with Figure S3c-d). Yet importantly, the performance of COMSV was only minimally reduced when it was given the real, noisy alignments as compared to having the artificial, correct alignments, showing that it does not require alignments of very high quality as input. This is important since for real cancer samples, molecule alignments are expected to contain a high level of errors.

Besides, even when the correct alignments were supplied to all methods, COMSV still performed substantially better than the other methods when calling indels from the samples with cancer sub-clones (S6 and S7) (Figure S3a-b), suggesting that these other methods had additional limitations unrelated to their reliance on correct alignments.

Next, we compared COMSV with the Bionano rare variant pipeline (RVP), which was specifically designed for detecting SVs from optical mapping data with a low fraction of molecule support. Since this pipeline was mainly designed for optical mapping data produced from DLS, we produced a new set of simulated data based on the recognition sites of the Direct Labeling Enzyme 1 (DLE-1). As seen in Figure 2f, COMSV outperformed RVP and the other three methods on the DLS data, especially in detecting SVs specific to the cancer sub-clones. RVP performed better than the original Bionano Solve in detecting rare inversions and duplications specific to the cancer sub-clones, but it still failed to detect many insertions and deletions in sub-clone 2.

Overall, these results show that COMSV is able to detect SVs of various types with high precision and sensitivity, from optical mapping data labeled by either a nicking enzyme or DLS. The advantage of COMSV over the other existing methods is most prominent when detecting SVs supported by few molecules such as those contained only in specific cancer sub-clones.

### 2.3 Identification of SVs from cancer samples

With the ability of COMSV in identifying SVs from synthetic data with cancer properties verified, next we applied it to identify SVs from real cancer samples. We obtained optical mapping data, including previously published data and some data newly generated in this study, from cancer and non-cancer cell lines and patient samples (Table S2). By comparing the SVs identified from the cancer and non-cancer samples, especially matched blood samples from the same patients, cancer-specific SVs could be identified. The matched non-cancer samples also helped evaluate the sensitivity of our SV calls, because most SVs identified from the non-cancer sample should also be detected in the corresponding cancer sample. Our pipelines support data produced by either nicking enzymes or DLS (Methods).

From each cancer sample, we detected 865-2,525 SVs (Figure 3, Table S3). We verified the reliability of these SVs using two different methods. First, we checked whether the SVs have been previously reported in the general populations, in cancer samples, and in exactly the same samples but having the SVs detected using a non-optical mapping method (Figure S4, Methods). From the results (Figure 3, Table S3), we found that 53.5%-92.9% of the SVs were previously reported in non-cancer samples from general human populations. These SVs are therefore likely not cancer-specific but are simply alleles not contained in the reference genome. Among the remaining SVs, some were previously reported in cancer samples by the Pan-Cancer Analysis of Whole Genomes (PCAWG) consortium, including SVs identified only from the same cancer/tissue type of our samples and those that were also identified from other cancer/tissue types. These SVs constituted up to 14.1% of all SVs we found in a sample. For some of our samples, we also had SVs previously identified using other types of data, including short-read sequencing, linked-read sequencing, and Hi-C. Overall, on average ∼80% of the SVs we identified from each sample have direct support from at least one of these three types of previous studies. In contrast, when we took the SVs identified by COMSV from all the DLE-1 and Nt.BspQI data sets and moved them to random locations in the genome, only 22.1% and 35.9% overlapped with these previously reported SVs, respectively (Table S3), showing that the fraction of SVs identified by COMSV from the samples having independent support is much higher than this count- and size-matched random SV set. On the other hand, we have also identified a large number of novel SVs, at an average of 375 of them per sample. As a community resource, SVs identified from individual samples are provided in Supplementary File 1 and a unified non-redundant list of all the novel SVs is provided in Supplementary File 2. These novel SVs were likely missed by the previous studies because of their cancer-type specificity (some cancer types of our sample are missing in PCAWG, such as nasopharyngeal carcinoma), sample specificity, and difficulties in identifying them without the long-range information provided by optical maps.

**Figure 3:**
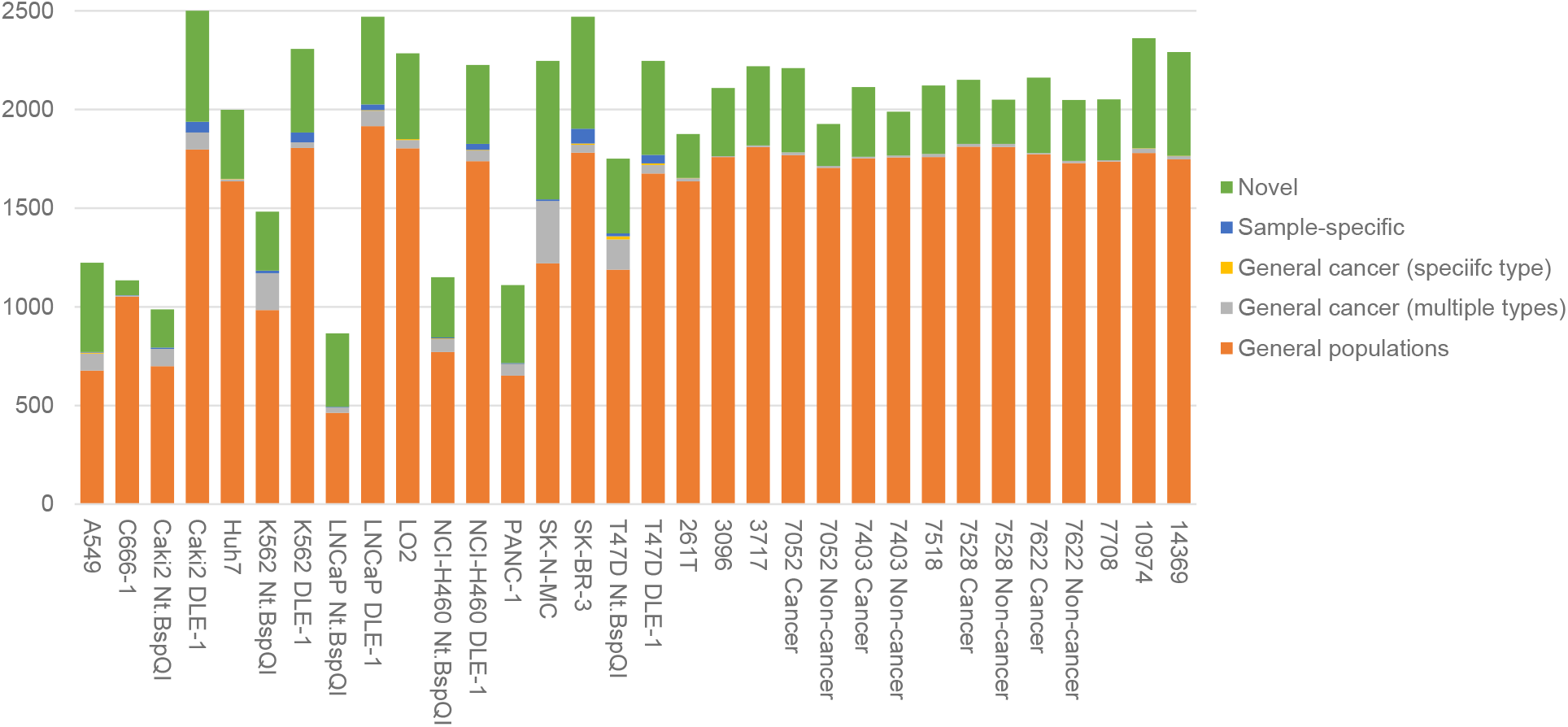
SVs identified by COMSV from the real samples. For each set of data, the identified SVs are put into different categories based on supporting evidence of them from independent data.

Next, we compared the SVs identified from matched cancer and non-cancer samples obtained from the same patients. For the four tongue squamous cell carcinoma samples, optical mapping data from blood samples of the same patients were also produced. Calling SVs from each sample independently and then comparing these SV call sets (Table S4), we found that the call sets of the tumor and blood samples from each patient are more similar to each other than to either cancer or blood call sets from other patients. This observation is consistent with the large proportion of non-cancer-specific SVs we found from the samples, as discussed above. Furthermore, in all four pairs of samples, we found that the proportion of SVs called from the blood samples that were also called from the matched cancer samples (86.8%-90.0%) is larger than the proportion of SVs called from the cancer samples that were also from the matched blood samples (77.2%-85.3%) (Table S4), which is in line with the expectation that the cancer samples contain somatic SVs not present in the matched blood samples.

To illustrate the different types of SVs identified by COMSV from the cancer samples, we visualize the alignments involved in several examples. Figure 4 shows two examples of insertions called from the cancer samples. Figure 4a shows a 2.6kb insertion on chromosome 19. This insertion was called in five samples, including 7403 Cancer, 10974, Caki2, K562, and LNCaP. From the molecule alignments in 7403 Cancer and K562 (Figure 4a), it is clear that the insertion is heterozygous in both samples, although the fraction of molecules that support the insertion is closer to 50% in 7403 Cancer than in K562 (which is <40%). In contrast, in samples 7052 Cancer and 7528 Cancer from which the insertion was not called, all molecules aligning to this locus do not support the presence of an insertion. Histograms of the distance between the anchoring labels also clearly show a bimodal distribution of distances in 7403 Cancer and K562, but a unimodal distribution of distances close to the expected distance according to the reference genome in 7052 Cancer and 7528 Cancer. Based on the two anchoring labels between which the distance on the supporting molecules is larger than the expected distance on the reference, the insertion enclosing region (i.e., the region within which the insertion happens) overlaps the *ZNF429* gene. *ZNF429* was reported to have the highest mutation frequency (36% with missense mutations) in the rare thymoma and thymic carcinoma in a small study of 14 samples [26], but otherwise there were few former reports of mutations of this gene in cancer. Our finding that this SV is contained in five cancer types from different tissues (tongue squamous cell carcinoma, hepatocellular carcinoma, kidney clear cell carcinoma, chronic myelogenous leukemia, and prostate adenocarcinoma) may indicate a previously unappreciated role of this novel SV and the *ZNF429* gene in cancer.

**Figure 4:**
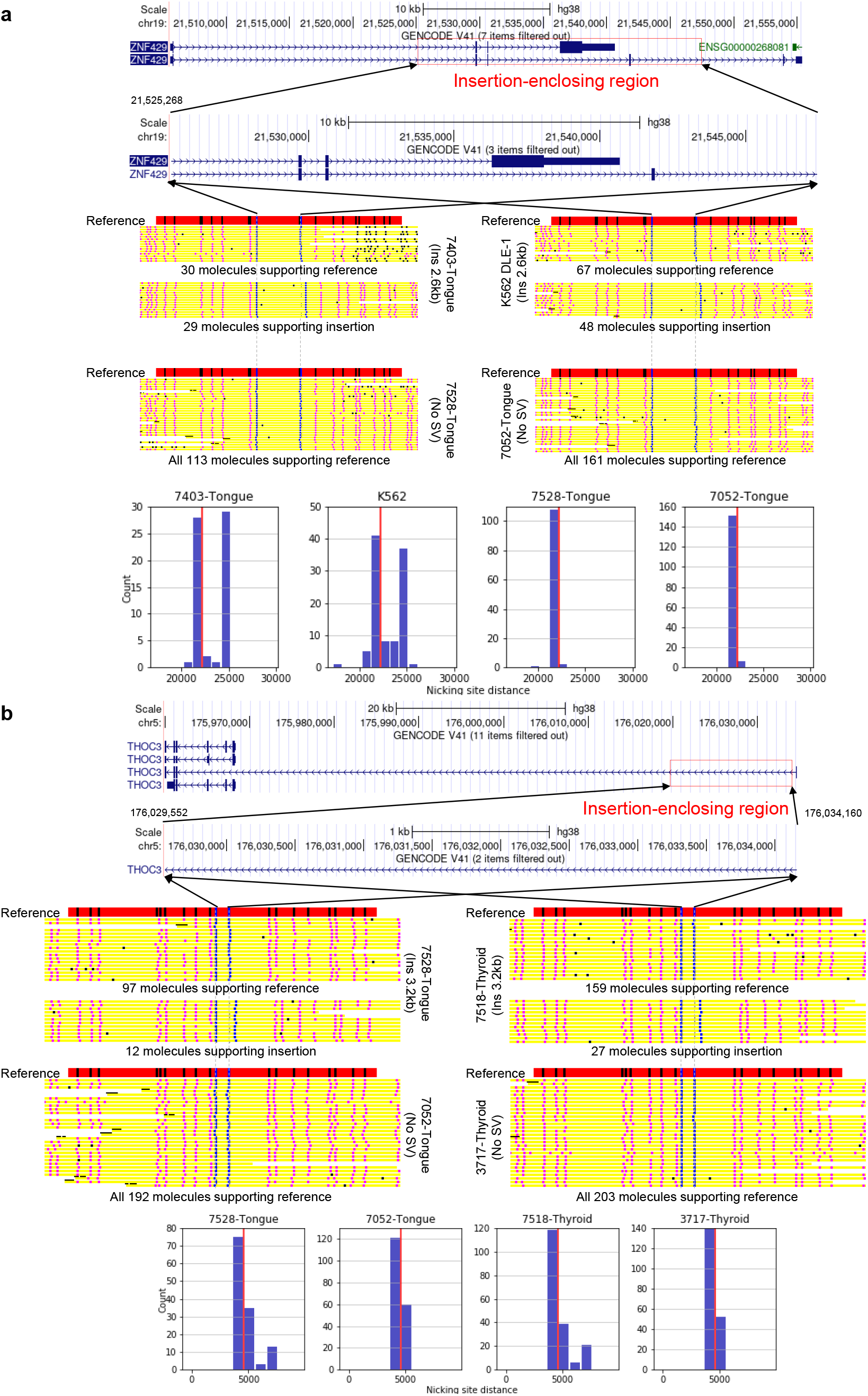
Examples of insertions called, from chromosome 19 (a) and chromosome 5 (b). In each panel, the top part shows the detected location of the insertion and a gene that it overlaps. The middle part shows the expected label pattern in the locus and the observed label patterns on optical maps from samples that have the insertion called or not called. A blue, pink and black dot on a molecule corresponds to an anchor label, a non-anchor aligned label, and an unaligned label, respectively. If the number of molecules aligning to the locus is too large, only the molecules with the largest and smallest distances between the anchoring labels are shown. The lower part shows the observed distance between the two anchoring labels between which the insertion is called. The expected distance according to the reference genome is marked with a red vertical line.

Similarly, Figure 4b shows a 3.2kb insertion on chromosome 5 that is identified in four samples, including 7518 Cancer, 7528 Cancer, LNCaP, and SK-BR-3. Again, the insertion is found to be heterozygous in 7518 Cancer and 7528 Cancer, but only 27/186=15% and 12/109=11% of the aligned molecules support the insertion in these samples, respectively. Considering that a) the absolute numbers of supporting molecules (27 and 12) are still fairly large, b) the quality of these alignments is good, as judged by the consistency of their label patterns and the expected pattern on the reference genome, and c) the insertion was called independently in four samples, the insertion calls should be reliable. This example illustrates COMSV’s ability to identify SVs even when the fraction of SV-supporting molecules is very low.

Figure 5 shows two examples of deletions called from the cancer samples. Figure 5a shows a 3.7kb deletion on chromosome 8. The deletion was called from two samples, including 7052 Cancer (tongue squamous cell carcinoma) and 10974 (hepatocellular carcinoma). In both samples, there is a clear bimodal distribution of distances between the two anchoring labels, although the fraction of molecules supporting the deletion is lower in 7052 Cancer (56/114=49%) than 10974 (66/87=76%) (Figure 5a). The deletion-enclosing region overlaps the gene *CSMD1*, which is a tumor suppressor gene in breast cancer [27] and esophageal squamous cell carcinoma [28].

**Figure 5:**
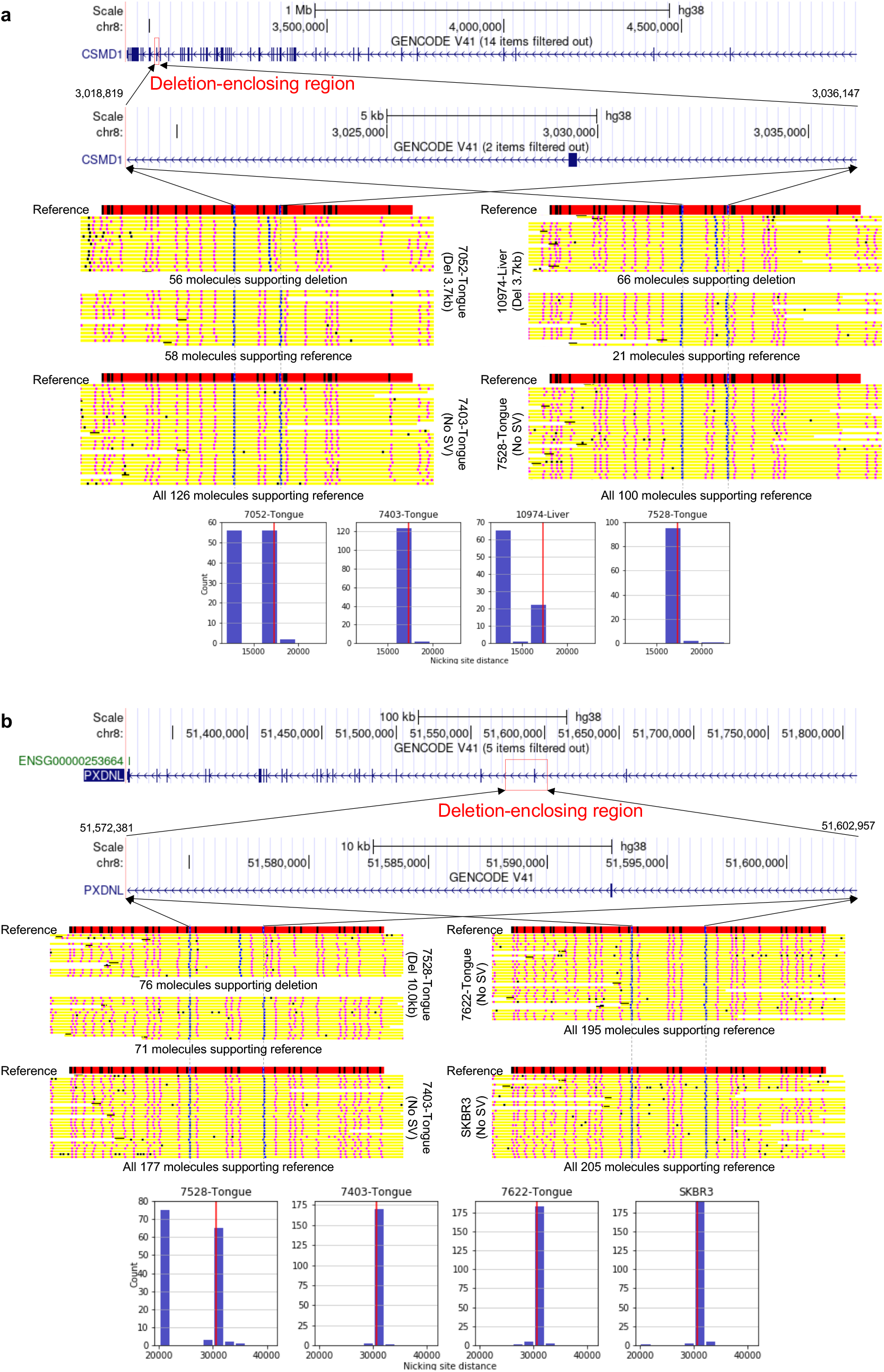
Examples of deletions called, both from chromosome 8. In each panel, the top part shows the detected location of the deletion and a gene that it overlaps. The middle part shows the expected label pattern in the locus and the observed label patterns on optical maps from samples that have the insertion called or not called. A blue, pink and black dot on a molecule corresponds to an anchor label, a non-anchor aligned label, and an unaligned label, respectively. If the number of molecules aligning to the locus is too large, only the molecules with the largest and smallest distances between the anchoring labels are shown. The lower part shows the observed distance between the two anchoring labels between which the deletion is called. The expected distance according to the reference genome is marked with a red vertical line.

Figure 5b shows a large, 10.0kb deletion on chromosome 8 that was found only in sample 7528 Cancer. The molecule alignments clearly show that the deletion is heterozygous in this sample.

Finally, Figure 6 shows four examples of complex SVs called from the cancer samples. Figure 6a shows an inversion identified on chromosome 8 from the thyroid anaplastic carcinoma sample 3717. 102/152=67% of the aligned molecules support the reference allele, while the remaining 50/152=33% of the aligned molecules support an inversion. These inversion-supporting molecules align well to a contig assembled *de novo* from the optical maps, which clearly shows an inverted pattern at the locus. This inversion is also identified from the bladder cancer sample 3096.

**Figure 6:**
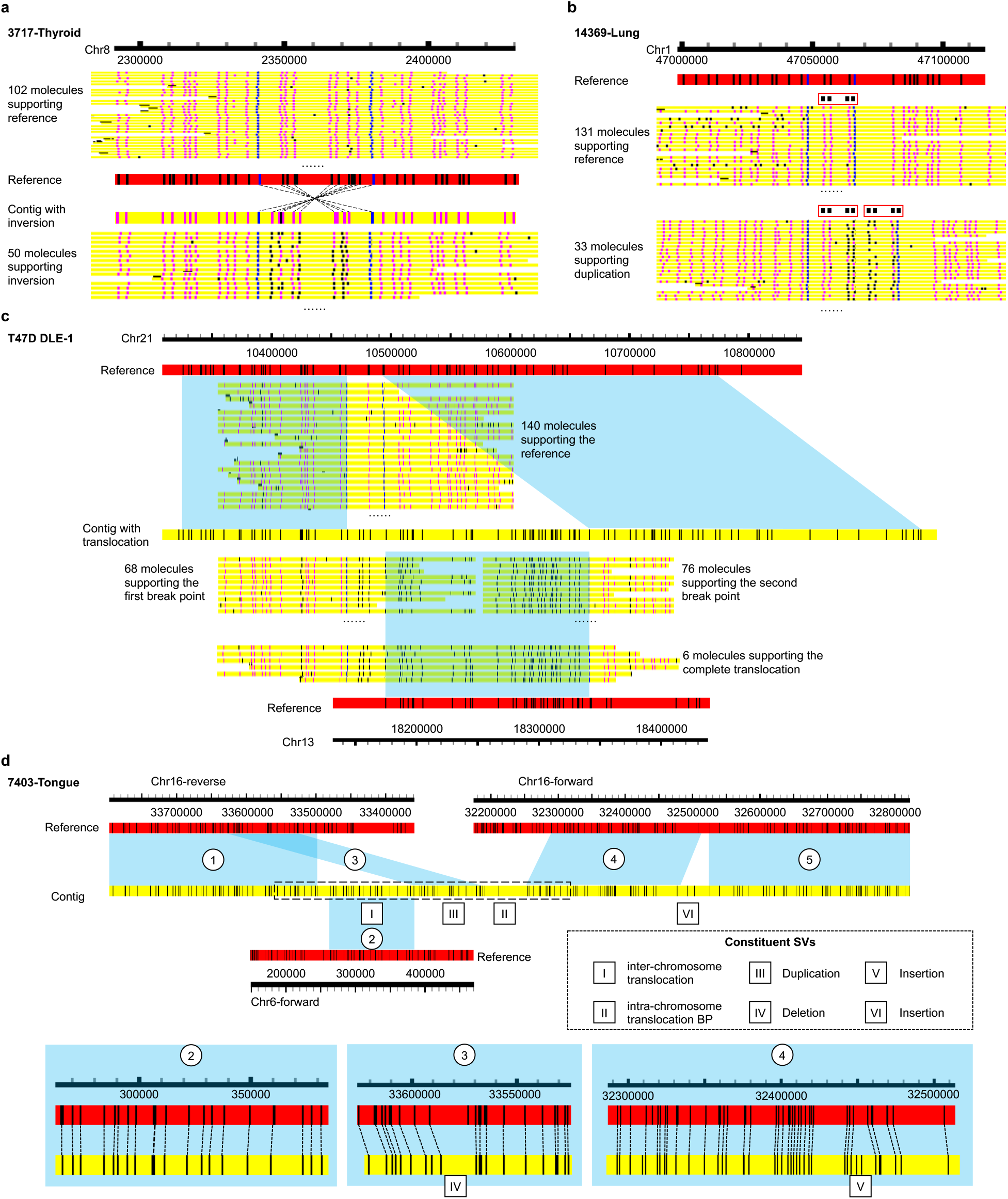
Examples of complex SVs called. **a** An inversion called on chromosome 8. In addition to the individual molecules, the contig assembled from the molecules and its alignment with the reference are also shown. **b** A tandem duplication called on chromosome 1. Each copy of the duplicating unit is highlighted by a red rectangle. **c** An inter-chromosomal translocation called on chromosomes 21 and 13. Segments of a contig and individual molecules aligned to the reference are indicated by blue parallelograms. In all three panels, a blue, pink and black dot on a molecule corresponds to an anchor label, a non-anchor aligned label, and an unaligned label, respectively. If the number of molecules aligning to the locus is too large, only a sample of the molecules is shown. **d** A very complex set of SVs called on chromosomes 16 and 6. Aligned segments of the contig are indicated by blue parallelograms and numbered (1-5). The individual constituent SVs are indicated by roman numerals (I-VI). The box with dashed line boundaries indicates the region with molecule alignments shown in Figure S5.

Figure 6b shows an example of tandem duplication identified on chromosome 1 from the lung pleomorphic carcinoma sample 14369. 131/164=79.9% of the aligned molecules support only one copy of the repeating unit, defined by a pattern of four labels. The remaining 33/164=20.1% of the aligned molecules show a second tandem copy of the repeating unit right next to the first copy.

Figure 6c shows an example of inter-chromosomal translocation identified from the breast ductal carcinoma cell line T47D. A contig assembled *de novo* from the molecules has a left segment and a right segment aligned to adjacent loci on chromosome 21, while the middle segment between them is aligned to chromosome 13. There are 140 molecules whose alignments to the reference span the two loci without any split alignment, thus supporting the reference allele. On the other hand, there are 68 molecules that are split-aligned to the first segment on chromosome 21 and to chromosome 13, which support the first set of translocation break points. Similarly, there are 76 molecules split-aligned to the second segment on chromosome 21 and to chromosome 13, which support the second set of translocation break points. Finally, there are 6 molecules whose alignments cover the first segment on chromosome 21, followed by chromosome 13, followed by the second segment on chromosome 21, and therefore they support the whole translocation. Interestingly, this translocation (or some of its break points) is also identified from 15 other samples (Huh7, LNCaP DLE-1, LO2, H460 DLE-1, 261T, 3096, 3717, 7052 Cancer, 7403 Cancer, 7518, 7528 Cancer, 7622 Cancer, 7708, 10974, and 14369). The large number of samples from which the translocation signals are independently detected and the presence of molecules that support the whole translocation event make this a very reliable translocation call.

Finally, Figure 6d shows a very complex set of SVs identified from the tongue squamous cell carcinoma sample 7403 Cancer. The contig assembled *de novo* from the molecules is partly aligned to chromosome 16 and partly aligned to chromosome 6, which form an inter-chromosomal translocation (I). The two segments of the contig aligned to chromosome 16 are aligned to different regions of chromosome 16 in opposite orientations, thus forming an intra-chromosomal translocation (II). The first segment shows patterns of a duplication (III), with a duplicated copy containing a deletion (IV). The other segment shows patterns of two insertions (V and VI). Since the reliability of this complex set of SVs depends on the correctness of the contig, we checked the alignments of individual molecules to it and found that it is supported by a large number of aligned molecules (Figure S5).

Overall, these examples demonstrate the ability of COMSV in calling SVs of different types and zygosities, including those with only a small proportion of supporting molecules.

## 3 Discussion

Despite recent advances in long-read sequencing technologies and the release of new high-throughput platforms (such as the PacBio Revio system), optical DNA mapping still has the longest average read length of over 300kb from routine procedures. The cost of optical DNA mapping is also relatively low (US$500 for 500× coverage of the human genome) as compared to long-read sequencing, making it a preferred platform for detecting large structural variations. Together with low-coverage sequencing data, the precise break points at base resolution could also be determined.

For the SVs identified by COMSV from each of the human samples, we found supports from previous studies for an average of 80.0% of them (ranging from 57.0%-93.3%). These SVs were previously identified from the general population, cancer samples (specific to the same cancer/tissye type or also other cancer/tissue types), and the same samples (based on non-optical mapping methods). One limitation of this analysis is the uneven coverage of the samples in previous studies. Whereas some cancer types were extensively studied, some others (e.g., nasopharyngeal carcinoma) were missing from large-scale cancer studies such as PCAWG. In addition, some particular samples had SVs called using multiple types of experimental methods previously, while other samples had none. Despite these differences, in each sample at least 6.7% of the SVs identified by COMSV were not found in the previous studies considered. We believe this is due to a combination of both the use of optical DNA mapping, which provides long read length, and the higher sensitivity of COMSV as compared to other existing SV calling methods. It would be useful to further validate these SVs and investigate their functional significance, especially the ones that COMSV independently identified from multiple samples.

For the samples with both Nt.BspQI and DLE-1 data, the two sets of SVs overlapped but each also contained unique SVs. This could be due to a number of reasons. First, DLE-1 in general has a higher density of labels than Nt.BspQI, allowing it to provide more information for calling SVs in general. However, there are also some loci at which Nt.BspQI produces more labels than DLE-1 (Figure S6). Second, DLE-1 prevents double-strand breakage at “fragile sites” caused by nearby nicks on opposite strands, leading to longer DNA molecules and thus better preservation of long-range information for SV calling. Third, due to different labeling densities and patterns, alignment and assembly methods optimized for one labeling method may not work well, or not without careful re-calibration, for the other labeling method. Alignment/assembly errors thus caused may have led to some of the inconsistencies in the two sets of SVs. Finally, although the optical mapping data sets involved in this study had high depth of coverage in general, some particular loci have lower coverage in the data for one labeling method than the other, which affects the ability to detect SVs at these loci, especially for the SVs with low allele ratio caused by sub-clones or cell type mixture.

## 4 Conclusions

In this study, we have proposed the COMSV method for identifying structural variations based on optical DNA mapping data produced from cancer samples, which is difficult due to cell type mixture, presence of sub-clones, aneuploidy and other copy number changes, large number of SVs, and complex SVs. We have shown that COMSV achieves good precision and sensitivity even if the SV allele is only observed in a small proportion of molecules aligned to the locus. Applying COMSV to cancer cell lines and patient samples, we have identified various novel SVs not reported in previous studies, including SVs that we recurrently but independently called from multiple samples. We have provided the SVs identified from the human samples, which cover a wide range of tissue and cancer types, as a community resource for studying cancer genomes.

## 5 Methods

### 5.1 Details of the indel pipeline

Before running COMSV pipeline to call SVs, optical mapping data of individual DNA molecules are aligned to a reference. For all the results in this study obtained from nickingenzyme-based optical mapping data, molecules were aligned using OMBlast [29] (in OMTools [30] v1.4a). For the results obtained from DLS, molecules were aligned using both OMBlast and RefAligner [19] (in Bionano Solve v3.5.1), with additional details given below. In both cases, in the alignment result, each molecule is either (fully or partially) aligned to a single genomic location, split-aligned with different parts aligned to different locations, or unaligned. For each aligned or split-aligned molecule, each label is either aligned to exactly one label on the reference or unaligned.

For each aligned molecule, COMSV first corrects for potential scaling errors as follows (Figure 1a). For every two adjacent labels on the molecule, the distance between them on the molecule divided by the distance between the two corresponding aligned labels on the reference is referred to as the observed-to-expected distance ratio. Each of these ratios is then divided by the median of all the observed-to-expected distance ratios from this alignment. For a perfectly linearized DNA molecule with all the labels correctly aligned, all these ratios should be close to one, with the small deviations caused by measurement errors alone. For a DNA molecule with imperfect linearization but a constant scaling factor along the whole molecule, after this normalization, all the ratios should also be close to one. Therefore, in both cases, if after the normalization there is a high proportion (*> a*%) of distance ratios strongly deviated from one (< 1 *− γ* or *>* 1 + *γ*), where *a* and *γ* are user parameters, the alignment is deemed low confident and marked for potential rejection (the whole list of parameters of COMSV and their default values are given in Table S5).

Whereas some of these low-confidence alignments can be incorrect alignments, some others can have abnormal distance ratios because of the presence of SVs instead. To distinguish between these two cases, the aligned genomic locations of these low-confidence molecules are analyzed. Specifically, all low-confidence alignments with overlapping aligned locations are grouped. For groups with more than *N* alignments, they are treated as reinforcing each other, considering the fact that the alignment of each molecule was performed independently. For such reinforced cases, all the alignments in the group will be “rescued”, while the other low-confidence alignments will be discarded.

The second step (Figure 1b) aims at collecting observed-to-expected distance ratios for the identification of indels in the later steps. Since indels can have different sizes and can occur at different locations, it would be ideal to check the distance between all pairs of labels no matter they are adjacent or not. However, computationally this is not feasible since it would require checking many label pairs. Instead, COMSV first starts with all pairs of labels that are adjacent either on the reference or on at least one molecule. For each of these label pairs, among all molecules containing both labels (according to their alignments to the reference), if at least *p_d_* of them have an observed-to-expected distance ratio strongly deviating from one (larger than 1+*δ* or smaller than 1-*δ*, where *δ* is the minimum SV ratio) and the absolute differences are larger than *s_min_* (minimum SV size), a suspected indel case is called. All these suspected indel regions are collected and the overlapping ones are grouped. Finally, for each group, all the label pairs within this region, no matter adjacent or not, will be considered in the calculation of distance ratios.

In the third step (Figure 1c), for each pair of labels collected in the second step, their distances on the molecules are clustered using complete-link hierarchical clustering until no two distance values respectively from any two clusters have an absolute difference smaller than *s_min_* or a ratio smaller than *δ*. Complete-link clustering is used because it is most sensitive to large-value outliers. The resulting clusters are tight, and outliers can be easily identified. When this clustering is finished, clusters that contain less than *n_s_* member molecules are considered outliers and are removed. The remaining clusters go through one more round of clustering using median-link hierarchical clustering (i.e., distance between two clusters defined as difference between their medians), which considers the opinions of all members in a cluster more evenly, to determine the number of clusters. Each final cluster is expected to contain molecules that contain a specific allele of the genomic region involved.

In the fourth step (Figure 1d), the clustering result for each pair of labels is first used to determine the SV type. Specifically, based on the number of clusters *n_c_*and the average observed-to-expected distance ratio *s_i_* of each cluster *i*, the following rules are used to determine the SV type:

| Number of clusters | Average distance ratios | Conclusion |
| --- | --- | --- |
| $n_c = 1$ | $s_1 \gg 1$ | Homozygous insertion |
| $n_c = 1$ | $s_1 \approx 1$ | No SV |
| $n_c = 1$ | $s_1 \ll 1$ | Homozygous deletion |
| $n_c = 2$ | $s_1 \gg 1$ and $s_2 \gg 1$ | Multi-allelic double insertion |
| $n_c = 2$ | $s_1 \gg 1$ and $s_2 \approx 1$ | Heterozygous insertion |
| $n_c = 2$ | $s_1 \gg 1$ and $s_2 \ll 1$ | Multi-allelic insertion-deletion |
| $n_c = 2$ | $s_1 \approx 1$ and $s_2 \ll 1$ | Heterozygous deletion |
| $n_c = 2$ | $s_1 \ll 1$ and $s_2 \ll 1$ | Multi-allelic double deletion |
| $n_c > 2$ | | Complex SV |

In these rules, a distance is considered significantly smaller than («) or larger than (») a threshold if the normalized deviation is larger than *s_min_* in the respective direction.

All these initial indel-spanning regions are then grouped to identify redundant cases. Specifically, all the indel-spanning regions are sorted according to their starting locations. All regions that have a reciprocal overlap of *>*50% are linked, and finally all regions directly or indirectly linked are grouped together. From each group, one final indel is produced with its type and zygosity determined by a simple majority vote of all the initial indels, and its span determined by following the indel candidate with the highest coverage (number of aligned molecules covering the region) among all the candidates in the group. Final indels with a coverage less than *c* are considered low-confidence and are discarded.

### 5.2 Confidence scores of detected indels

We then trained a support vector machine model with a radial basis function (RBF) kernel for distinguishing the correct indel calls of COMSV from the incorrect ones. Specifically, for each indel identified by COMSV from the simulation data, we put it into one of three possible classes by comparing it with the ground truth, namely wrong call (the indel does not actually exist), correct call of indel location but wrong zygosity, and correct call of indel location and zygosity. For each indel, three features were derived based on the outputs of COMSV, namely the ratio of molecules that support the SV allele, the detected indel size, and the number of molecules aligned to the locus. All features were transformed to have a value range between 0 and 1, where this was done by subtracting the minimum followed by dividing by the maximum. Based on a class-stratified five-fold cross-validation, the trained model achieved an area under the receiver-operator characteristic (AUROC) of 0.98 in distinguishing between true and false indels (without considering zygosity). In contrast, models that involved single features alone achieved an AUROC of 0.75, 0.28, and 0.81, with the ratio of molecules that support the SV allele, SV size, and the number of molecules aligned to the locus, respectively. Since the model has a prediction score for each of the three classes, these scores can also be used to check whether it is more likely for an SV to be homozygous or heterozygous.

### 5.3 Details of the complex SV pipeline

The complex SV pipeline of COMSV uses split-alignments of molecules and partially unaligned molecules to identify SVs. In a split-alignment, two or more segments of a molecule are aligned to the reference non-consecutively. Depending on where the different segments are aligned and the orientations of these individual alignments, different types of SVs are called (Figure 1e). In general, if a molecule can provide information about one break point of an SV but not the complete information of the whole SV, only an SV break point is called.

The exact way that SV break points and complete SVs are called depends on the SV type (Figure 1e). For a translocation, there should be three break points in total for a complete event, namely two break points between which a DNA segment is cut, and a third break point at which the segment is inserted. If a molecule contains two segments split-aligned to two regions on the same chromosome separated by at least 5Mb, two intra-chromosomal translocation break points (marked as “-BP” in the final SV lists) are called. Specifically, suppose a segment is aligned to region [*A, B*] and a later segment is aligned to region [*C, D*] in the same alignment orientation, with *A < B < C < D*, then two break points are called between *B* and *C*, with one expected to be close to *B* and the other one expected to be close to *C*. The actual translocation can involve either a segment that starts near *C* and ends at an unknown location after *D* translocated to a location near *B*, or a segment that starts at an unknown location before *A* and ends near *B* translocated to a location near *C*. Now, suppose after the second segment, there is a third segment on the molecule aligned to the region [*E, F*], with *B < E < F < C*, then a complete translocation event is reported with a segment that starts near *C* and ends near *D* translocated to a location between *B* and *E*. A similar logic can be used to call a complete intra-chromosomal translocation if the third segment is before the first segment rather than after the second segment. Interchromosomal translocations can also be called in a similar way, except that the translocated segment should come from a chromosome different from the one at which it is inserted.

For an inversion, there should be two break points in total for a complete event, between which the DNA is inverted. Suppose a segment is aligned to region [*G, H*] and a later segment is aligned to region [*I, J*] in the reverse alignment orientation, with *G < H < I < J*, then one inversion break point is called and its location is between *H* and *I*. The inversion should involve a segment that starts at this break point and ends at a location after *J*. Now, suppose after the second segment, there is a third segment on the molecule aligned to the region [*K, L*], with *K > J*, then a complete inversion is called with the inverted segment starting at a location between *H* and *I* and ending at a location between *J* and *K*. In some cases, the inverted segment has a symmetric pattern that makes it difficult for the aligner to detect an inversion using the split-alignment strategy. If there are clear distance changes of the labels flanking the inverted segment, it is possible for COMSV to detect an insertion and a deletion instead (Figure S7). To detect these cases, we collected the deletion-insertion pairs with genomic distance less than 20kb and size difference less than 5%. These pairs are re-considered as inversions if they concurrently appear in at least 2 molecules.

For a duplication, there should be three break points in total for a complete event, namely two break points between which a DNA segment is duplicated, and a third break point at which the segment is inserted. If a molecule contains two segments with their aligned regions on the reference overlapping each other, the starting and ending locations of their overlapped region are called as two duplication break points, where the segment duplicated should completely enclose this region. Other parts of the molecule can help refine these break points and identify the third break point. For example, suppose the overlapped region of the two alignments is [*O, P*], if the segment on the molecule between these two aligned segments is aligned to region [*M, N*] on the reference, where *M < N < O < P*, then the duplicated region may start somewhere between *N* and *O*. On the other hand, if a later segment of the molecule is aligned to region [*Q, R*] on the reference, where *Q* is much larger than *P*, then the duplicated DNA is expected to have inserted to a location near *Q*.

A special case of duplications is that the whole molecule is aligned to consecutive regions on the reference except that two adjacent segments are aligned to the same region. In this case, a tandem duplication is called.

Finally, if different segments of a molecule are aligned to nearby genomic regions within 5Mb, an insertion or deletion will be called depending on the aligned locations of the segments. Some of these indels may have already been identified by the indel pipeline, but some may only be identified by this complex SV pipeline especially in the case of a large deletion, where the different segments cannot be aligned to the reference without a split-alignment.

The complex SV pipeline of COMSV also makes use of partially aligned molecules to identify duplications and inversions. Specifically, for each molecule that is only partially aligned to the reference, the unaligned segment is extracted and aligned to the reference again by itself. It may be uniquely aligned to a single region on the reference or to multiple different regions. In the former case, if the new alignment has an orientation different from the original aligned segment, an inversion break point or a complete inversion would be called, and if the new alignment region overlaps the one of the original aligned segment, a duplication would be called. On the other hand, if the alignment is not unique to a single region and one of these aligned regions supports neither an inversion nor a duplication, no SVs would be called.

It should be noted that in the above descriptions, break points can only be identified up to the closest aligned label, and therefore calling a break point precisely requires both the presence of a label (e.g., a nicking site) near the break point and a correct alignment of the label on at least one molecule.

If there are multiple complex SV break points that support overlapping SVs of the same type, they are grouped together to give one final break point call, with the possible range of the break point defined in three different ways, all reported in the output file. The three ways are: 1) taking the intersection of all the individual break point ranges, 2) taking the minimum span that covers all the individual break point ranges, and 3) taking the break point range with the highest molecule support. For example, for the inversion shown in Figure 1f, the molecules *m*_11_ and *m*_13_ propose the left break point to be within the regions *T − U* and *T − V*, respectively, and therefore in the final call, the common region of them, *T − U*, is taken as the range of the left break point.

For translocations and duplications, if the translocated/duplicated content is inverted when inserting to the new location, the SV type is recorded as “invert-translocation” and “invert-duplication”, respectively.

It is possible that some duplications are initially missed or identified as insertions if the duplicated parts are not aligned in the initial alignment. To deal with these cases, COMSV includes a module for “rescuing” unaligned segments of optical maps that have other parts properly aligned. Specifically, for each unaligned segment of an optical map, we check whether its label pattern matches any region on the reference near (<500kb) the aligned parts. If *≥*80% of the unaligned segment can match well, a duplication would be called. Similarly, if the reverse of the label pattern can match the reference of a nearby region well, an inversion would be called. All SVs identified by re-alignment of initially unaligned segments are marked as “-realigned” on the output SV list.

It is possible that some very complex types of SV appear in cancer genomes, such as nested SVs with one SV completely enclosing another SV. These cases can be identified for manual inspection and analysis by looking for regions with a high density of SVs identified by COMSV.

### 5.4 Customized methods for DLE-1 data

Compared to traditional optical mapping data using a nicking enzyme (such as Nt.BspQI) for labeling, DLS data using the DLE-1 enzyme for labeling have conceptually no fragile sites (i.e., sites easily having double-stranded breakage due to two nicks on opposite strands close to each other). As a result, in DLS data the molecules are longer on average. There is also a higher density of labels. These properties created difficulties for the alignment algorithms we used (especially OMBlast), which were originally developed for nicking enzyme-based data.

To deal with this issue, COMSV integrates SVs detected from multiple sets of alignments when working on DLS data, as follows.

Indel detection requires highly accurate alignment of individual labels. OMBlast was not optimized for DLS data and could produce alignments with high error rates. In contrast, RefAligner and Assembler in the Bionano Solve toolset have been optimized for DLS. Therefore, for DLS data, we used two sets of alignments produced by RefAligner in our indel pipeline, namely i) direct alignment of individual molecules to the reference, and ii) alignment of assembled contigs to the reference. Since the assembled contigs already represent a form of consensus among the individual molecules, when calling indels based on their alignments, we did not impose the requirements for molecule alignments (*N*, *n_s_*, and *c*). For an identified SV, if there was at least one contig that supported the reference allele, we called it a heterozygous SV. Otherwise, we called it a homozygous SV.

To identify complex SVs from DLS data, in addition to the two types of alignments used to detect indels, we also engaged OMBlast to do a two-round alignment of individual molecules, which is a strategy we also used in our previous studies [21, 24]. In the first round, molecules were aligned to the reference with the option of complex SV break points detection disabled, such that molecules that required only simple alignments could be performed reliably, while some molecules that required more complex alignments (such as those around SV break point regions) were either completely unaligned or had segments of them unaligned. In the second round, all unaligned molecules and molecule segments were aligned to the reference again, independent of the alignment results in the first round. The final complex SV callset was produced by applying our complex SV pipeline to all alignments (these two rounds of OMBlast alignment, RefAligner direct molecule alignment, and RefAligner contig alignment). These three sets of SVs were then integrated by including 1) all complex SVs detected from the RefAligner contig alignments, 2) all duplications detected from RefAligner direct molecule alignments (because RefAligner in Bionano Solve could not generate split alignments and thus could not be used to detect SVs such as inversions), and 3) SVs detected from OMBlast two-round split alignments of individual molecules, except for those that required the rescue procedure for unaligned segments as described above.

To improve the accuracy of our callset, we removed indels that overlapped any duplications or inversions detected by the complex SV pipeline, because the confidence of local alignments around complex SV breakpoints was generally lower and the indel pipeline was more likely to generate false positive calls.

### 5.5 Defining the overlap between an SV and another SV or a genomic region

In various parts of this study, we had to compare the location information of an SV with that of a simulated SV, an SV reported in a previous study, or an annotated gene.

To do that, we first needed to define the “spanning region” (or simply “span”) of an SV, which is basically where the SV appears. Conceptually, the span of an insertion is the single nucleotide position at which the extra DNA sequence is inserted, the span of a deletion is the genomic region deleted, the span of an inversion is the genomic region inverted, and the span of a duplication is the genomic region duplicated. In a translocation, a genomic region is translocated to another region, which involves a deletion and an insertion, and thus it has two spans. Among the two spans of a translocation, the one with a smaller starting point (earlier chromosome, based on the order of 1, 2, …, 22, X, Y, or same chromosome but smaller coordinate) is called the first span and the other one is called the second span.

In all these cases, due to the nature of optical mapping, a span of an SV can only be estimated based on the innermost pair of labels that enclose it. The region between these two labels on the reference was the practical definition of a span used by COMSV.

Accordingly, we defined different types of overlaps that involve an SV as follows:

- For non-translocations, two SVs of the same type are considered overlapping if their spans have reciprocal overlaps of *≥*50%.
- For non-translocations, an SV is considered overlapping with a genomic region (e.g., a gene or a gold-standard SV, which has a precise span rather than an estimated one) if the span of the SV has an overlap with the genomic region of *≥*1bp.
- Two translocations are considered overlapping if the shortest distance between their first span is *≤*500kb and the shortest distance between their second span is *≤*500kb.
- A translocation is considered overlapping with a genomic region if the shortest distance between the genomic region and either span of the translocation is *≤*50kb.

In comparisons involving SV locations, we sometimes also considered additional requirements, such as the size of an SV. For example, even if two insertions have the same span (i.e., estimated insertion point), they would be different insertions if the lengths of their inserted sequences are very different. For parts of our study that introduced such additional requirements, the details are provided in the corresponding method descriptions separately.

### 5.6 Procedures for generating simulated data

In the first simulation experiment, we generated large indels with the proportion of supporting molecules ranging from 10% to 90% as follows. First, following the parameter settings of a previous work [21], we simulated a list of indels on the hg38 human reference genome using pIRS [31]. We then took the reference genome as one haplotype and the indel-added genome as another haplotype to simulate molecules using *in silico* digestion to 100x average genome coverage from each haplotype, using the same parameter settings as in a previous work [21]. Finally, molecules from these two sets were randomly sampled at different rates to produce 17 data sets, respectively corresponding to sampling 10%-90% of the molecules from the reference haplotype and the remaining from the indel-added haplotype, with an increment of 5% each time.

In the second simulation experiment, we first generated the genomes of normal cells, trunk cancer cells, and cells in the two sub-clones by simulating different sets of SVs using pIRS based on the evolution graph (Figure S2). The actual numbers of SVs generated are shown in Table S6. We then simulated molecules for each genome to 100x average coverage. These molecules were then sampled to form the molecules of the seven samples according to their target cell compositions (Table S1).

The above data sets were simulated based on the Bionano Irys platform (which uses the Nt.BspQI enzyme for labeling). To check the performance of COMSV on data produced by the Saphyr platform (which uses the DLE-1 enzyme for labeling), we also generated simulated optical mapping data by applying DLE-1 *in silico* digestion to molecules of Sample 7 in Table S1.

### 5.7 Application of the different SV callers on the simulated data

We compared COMSV with four other SV callers on the simulated data, namely the two SV callers included in Bionano Solve v3.5.1 (regular pipeline and RVP), OMIndel, and OMSV. For the regular Bionano Solve pipeline, the exact list of command-line arguments used was “$BIONANOSOLVE DIR/Pipeline/1.0/pipelineCL.py -l prop${sample} -t $BIO-NANOSOLVE DIR/RefAligner/1.0 -b $BNXPATH/${sample} -y -d -U -i 5 -F 1 -W 1 -c 1 -a optArguments haplotype saphyr human.xml -r hg38 BSPQI 700.cmap -f 0.2 -J 48 -j 60 -jp 80 -T 80 -N 6”. We ran the RVP pipeline via the Bionano Access platform with the same set of arguments used in the regular pipeline. For OMIndel, we tried different values of the minimum supporting reads and picked the one that gave the highest F1 score across samples, which was found to be 7. Default values were used for all other parameters. When zygosity was considered, the minimum cluster size of the reference reads was set to 10, as suggested by the author. For OMSV, we set the minimum support to 10 and the minimum likelihood ratio to 1*e*4, which provided the best performance on the simulated data.

After obtaining the callsets of the different SV callers, we compared them with the gold-standard list of SVs after removing SVs that overlapped fragile sites, regions with no sequence information on the reference, and pseudo-autosomal regions. A called SV was considered correct if it overlapped with a gold-standard SV of the same type with the estimated SV size within 5 times of the correct size. For complex SVs, we also required each called break point to be within 20kb from a break point of the gold-standard SV. Precision, recall, and F1 score were then computed accordingly.

### 5.8 Collection of published optical mapping data

We collected published optical mapping data from several sources (Table S2). All published cell line data with DLE-1 labeling were from a previous study [32] and kindly provided to us by Vineet Bafna in the form of BNX files. The C666-1 data were produced in our previous work [21] and we used the alignment files from that study directly. All other published cell line data with Nt.BspQI labeling were produced in our previous work [10] and we took the BNX files as our input. The published patient sample data were from a previous study [33] and kindly provided to us by James Broach, with the help of Lijun Zhang, in the form of BNX files.

### 5.9 Production of optical mapping data from cell lines and primary samples

The two immortalized human liver cell lines, Huh7 and LO2, were maintained in Dulbecco’s modified Eagle’s medium (Gibco) with 10% FBS (HyClone). All cells were cultured at 37°C in a humidified chamber containing 5% CO2.

A HCC patient who underwent hepatectomy at the Prince of Wales Hospital (Hong Kong) was included in this study. The specimen was processed immediately after surgery and snapfrozen in liquid nitrogen for DNA extraction. The patient gave written consent on the use of specimen for research purposes. The study protocol was approved by the Joint Chinese University of Hong Kong-New Territories East Cluster Clinical Research Ethics Committee.

High molecular weight (HMW) genomic DNA was extracted from Huh7 and LO2 cells using Bionano Prep^TM^ Blood and Cell Culture DNA (Part no. 80004; Bionano Genomics, USA) with the Cell Culture Protocol (Document no. 30026 Rev F), and from 261T using Bionano Prep Animal Tissue DNA Isolation Kit (Part no. 80002) (Bionano Genomics, USA) with the Soft Tissue Protocol (Document no. 30077 Rev C). In detail, cell suspension was obtained by resuspending cell line pellets or homogenized solid tumor sample in 1X phosphate-buffered saline (PBS) buffer solution. Cell suspension was then embedded in low-melting agarose gel plugs and treated with protease K and RNase A. The gel piece was then molten with heat and agarose. HMW DNA was recovered from the molten gel via drop dialysis, and fluorescently labelled to generate sequence motif-specific patterns following the Bionano DLS protocol (Document no. 30206 Rev F). Optical mapping data were generated on the Bionano Saphyr system (Part no. 60325) using Saphyr Chip G1.2 (Part no. 20319). In brief, fluorescently labeled HMW DNA molecules were automatically stretched and imaged within nanochannel arrays. Distances between labels were calculated from the images into raw molecule maps and *de novo* assembled into consensus maps by Bionano Access v1.4.2 and Bionano Tools v1.4.1.

### 5.10 A non-redundant list of SVs identified from the real samples

For each real sample, COMSV produced a final list of SVs identified from it with various types of annotations, such as SV type, zygosity, ranges of genomic locations of the break points, and proportion of aligned molecules supporting the SV allele. Following our previous work [24], we excluded several classes of genomic regions from SV calling that make molecule alignment erroneous or even impossible (Supplementary File 3), including 1) regions with low density of Nt.BspQI or DLE-1 labels, defined as regions *≥*100kb with *≤*5 labels, 2) regions with unresolved sequences in the reference (“N” regions) *≥*50kb, 3) centromeres, and 4) pseudo-autosomal regions. We also excluded the Y chromosome.

To annotate the genes that overlap each SV, we downloaded the latest release (release 43) of comprehensive human gene annotation from GENCODE [34] in GTF format (https://ftp.ebi.ac.uk/pub/databases/gencode/Gencode_human/release_43/gencode.v43.annotation.gtf.gz). We considered only annotations on chromosomes 1-22, X, and Y. We annotated each SV by all the genes that it overlapped, including both coding genes and non-coding genes. For each gene, we listed the Ensembl gene ID, gene name, and gene type. For SVs that spanned *≥*5Mb, which may overlap many genes, we did not include their gene annotations.

Besides, we also integrated the SV lists from individual samples to produce an overall list of non-redundant SVs identified from all samples. To do that, we considered one SV type at a time. We collected all SVs of that type identified from all samples. For an SV type other than translocations, overlapping SVs (as defined above) were put into the same group. This was done in a transitive manner, such that if the first and second SVs were grouped, the second and third SVs were grouped, and so on, until the *k*-th and (*k* + 1)-th SVs were grouped, all (*k* + 1) SVs would all be grouped together. Each group was then represented by the SV in it that had the smallest genomic span. Due to this process of merging SVs across samples, in the final list of non-redundant SVs, we omitted annotations that required sample-specific information, such as zygosity and proportion of aligned molecules supporting the SV allele, but instead recorded whether a sample contained an original SV belonging to each group. We also simplified SV type annotations since the more specific types could be present in only some samples. For example, an “invert-duplication” (a duplication with the duplicated part inverted) was simply recorded as the more general “duplication” type.

Translocations were merged in a way slightly different from the other SV types due to their complexities. Conceptually, each translocation involves “cutting” a genomic region out and “pasting” it somewhere else. In practice, optical mapping data usually can only provide information about break points, by having different segments of a molecule aligned to different separated regions on the reference, without complete translocation information. Therefore, usually two different regions are identified to be involved in a translocation, but which one corresponds to the “cut” site and which to the “paste” site is not known. Because of this, when merging translocations called from different samples, both regions need to be considered. Two translocations were grouped only if both their first spans overlapped and their second spans overlapped. Procedure-wise, we sorted translocations based on their two spans’ genomic locations, followed by grouping translocations with both spans overlapping using the transitive rule described above. Finally, for each set of merged translocation break points, it was considered not called from a sample if there are *≤*4 molecules supporting it in the sample.

SV break points detected for which the complete SV could not be determined were marked as “-BP” in the output files.

### 5.11 Identifying SVs called by COMSV from the real samples that were reported in previous studies

We collected multiple sets of SVs from different sources to check whether the SVs called by COMSV from the real samples were also reported in some previous studies.

Two SV sets were from general populations. The first set included SVs larger than 2kb in size identified from 156 samples across 26 human populations using optical DNA mapping [24]. The second set was the NCBI Curated Common Structural Variants in dbVar [35], available at https://www.ncbi.nlm.nih.gov/dbvar/studies/nstd186/. These two sets together covered SVs of different sizes identified across various human populations.

SVs larger than 1kb in size previously identified from cancer samples were collected from the latest open Pan-Cancer Analysis of Whole Genomes (PCAWG) consensus callsets for structural variants [36], available at https://dcc.icgc.org/releases/PCAWG/consensus_sv. This database contained SVs from 2,588 cancer samples, each annotated with a cancer type. For samples with multiple SV files, we merged them into a single SV list for each sample. We omitted 190 samples that were not annotated with a cancer type.

Finally, for some specific samples, we collected previously reported SVs (from the supplementary materials of corresponding publications) larger than 1kb in size that were identified using non-optical mapping methods (Table S7). Again, for each sample with multiple lists of SVs, we merged them into a single list.

Based on these SV sets, we classified each SV identified by COMSV into one of five possible classes (Figure S4). If an SV was reported in either of the two general population SV sets (by having overlapping locations, the same type, and size ratio within [1/5, 5] with any known SV in the population sets for indels, or having the same type with any known SV as well as all breakpoints located within 10kb of that known SV in the population sets for complex SVs), we classified the SV as being found in the general population. Otherwise, we checked if the SV was reported by PCAWG. If it was, then we further checked whether it was reported in at least one cancer type other than the cancer type of our sample, in which case the SV was classified as being found in multiple cancer types, or else it was classified as being found in a specific cancer type. If the SV was not reported by PCAWG, then we further checked the SVs previously reported from this same sample, if available, to classify the SV as sample-specific (and previously reported) or novel.

## 6 Declarations

### 6.1 Ethics approval and consent to participate

A HCC patient who underwent hepatectomy at the Prince of Wales Hospital (Hong Kong) was included in this study. The patient gave written consent on the use of specimen for research purposes. The study protocol was approved by the Joint Chinese University of Hong Kong-New Territories East Cluster Clinical Research Ethics Committee.

### 6.2 Consent for publication

Not applicable

### 6.3 Availability of data and materials

The datasets used and/or analysed during the current study are available from the corresponding author on reasonable request.

### 6.6 Authors’ contributions

KYY conceived the study. LL and CH developed the methods. KWL, PBSL, JW, JZ and ASLC obtained and prepared the biological samples. JX, CYLC, TFC and FY produced the data. LL, CH, CYLC, AKYL, DB, LC, TFC and KYY analyzed the data. LL, CH and KYY prepared the manuscript. All authors read and approved the final version of the manuscript.

## 6.7 Acknowledgements

We thank Vineet Bafna, James Broach and Lijun Zhang for providing published optical mapping data. We acknowledge the assistance of Xian Fan and Ernest Lam in running OMIndel and Bionano Solve, respectively.

## S1 Supplementary materials

### S1 Supplementary tables

**Table S1:**
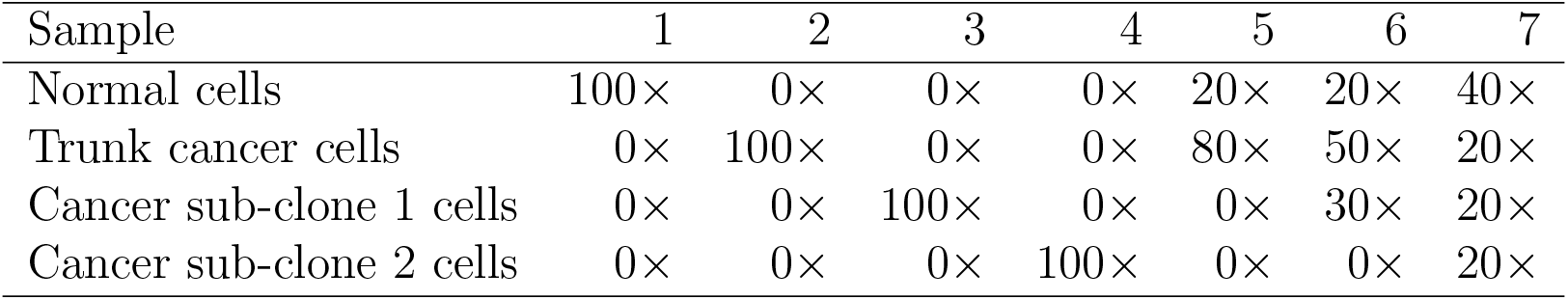
Cell type composition of the different samples in the second simulation experiment. For each sample, the amount of optical maps produced from each cell type in terms of average genome coverage is shown.

| Sample | 1 | 2 | 3 | 4 | 5 | 6 | 7 |
| --- | --- | --- | --- | --- | --- | --- | --- |
| Normal cells | 100× | 0× | 0× | 0× | 20× | 20× | 40× |
| Trunk cancer cells | 0× | 100× | 0× | 0× | 80× | 50× | 20× |
| Cancer sub-clone 1 cells | 0× | 0× | 100× | 0× | 0× | 30× | 20× |
| Cancer sub-clone 2 cells | 0× | 0× | 0× | 100× | 0× | 0× | 20× |

**Table S2:**
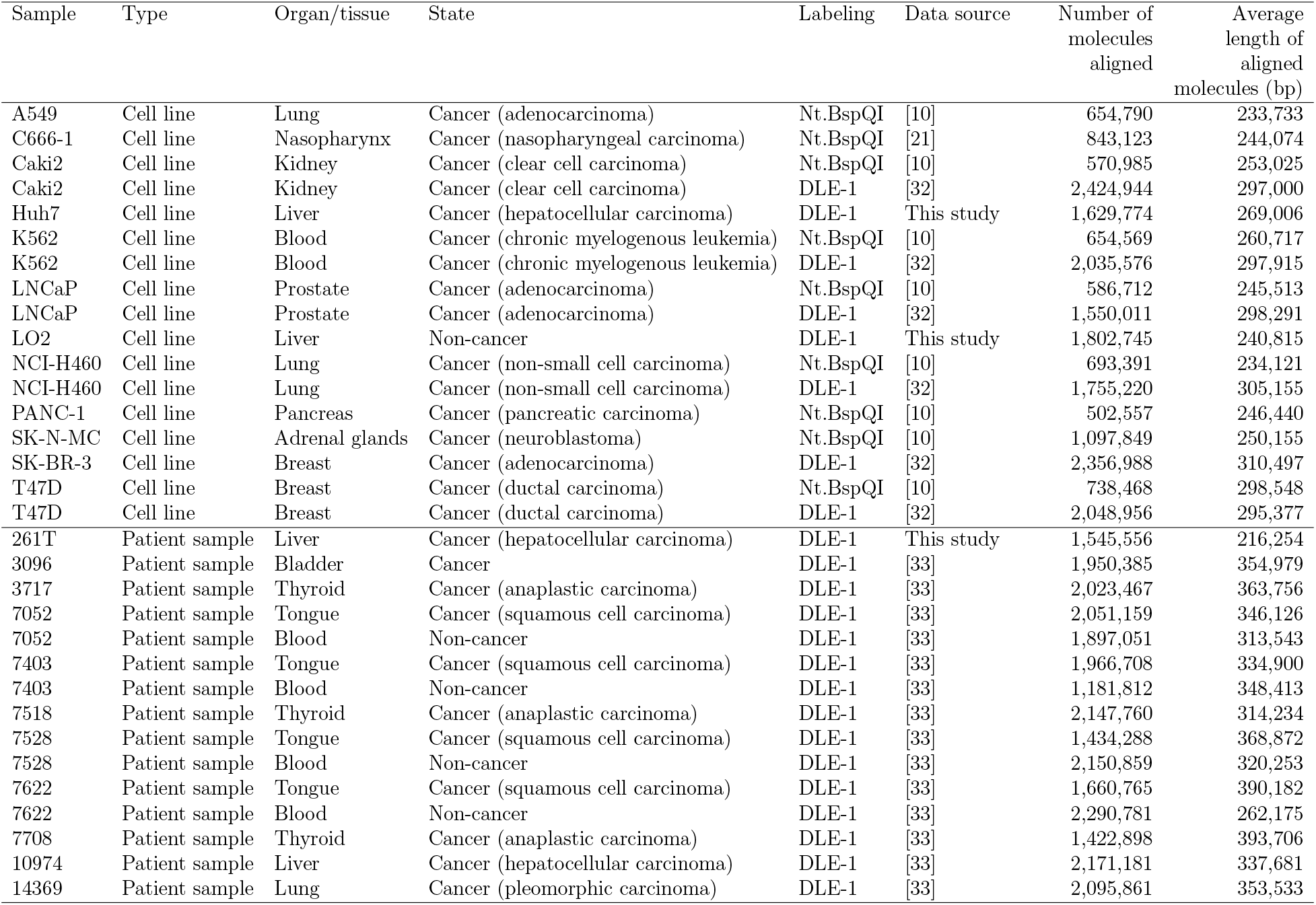
Optical mapping data of human samples included in this study.

**Table S3:** Numbers of SVs called from the human samples that overlapped with other SV call sets. For non-cancer samples, cancer-type-specific comparisons were performed based on the organ/tissue type. For samples without SV call sets previously obtained from non-OM data, their sample-specific overlap numbers are marked as “NA”. Translocations were excluded from the random sets because they cover two separate genomic spans instead of one.

| Sample | General pop-<br>ulations |  | General can-<br>cer (multiple<br>types) |  | General can-<br>cer (specific<br>type) |  | Sample-specific |  | Novel |  | Total |
| --- | --- | --- | --- | --- | --- | --- | --- | --- | --- | --- | --- |
| A549 | 677 | (55.36%) | 86 | (7.03%) | 5 | (0.41%) | 3 | (0.25%) | 452 | (36.96%) | 1,223 |
| C666-1 | 1,053 | (92.86%) | 4 | (0.35%) | 0 | (0%) | 1 | (0.09%) | 76 | (6.70%) | 1,134 |
| Caki2 Nt.BspQI | 699 | (70.89%) | 88 | (8.92%) | 0 | (0%) | 7 | (0.71%) | 192 | (19.47%) | 986 |
| Caki2 DLE-1 | 1,797 | (71.17%) | 86 | (3.41%) | 1 | (0.04%) | 53 | (2.10%) | 588 | (23.29%) | 2,525 |
| Huh7 | 1,635 | (81.83%) | 12 | (0.60%) | 1 | (0.05%) | NA |  | 350 | (17.52%) | 1,998 |
| K562 Nt.BspQI | 984 | (66.40%) | 186 | (12.55%) | 0 | (0%) | 14 | (0.94%) | 298 | (20.11%) | 1,482 |
| K562 DLE-1 | 1,805 | (78.24%) | 28 | (1.21%) | 0 | (0%) | 50 | (2.17%) | 424 | (18.38%) | 2,307 |
| LNCaP Nt.BspQI | 463 | (53.53%) | 27 | (3.12%) | 0 | (0%) | 3 | (0.35%) | 372 | (43.01%) | 865 |
| LNCaP DLE-1 | 1,916 | (77.57%) | 77 | (3.12%) | 3 | (0.12%) | 29 | (1.17%) | 445 | (18.02%) | 2,470 |
| LO2 | 1,803 | (78.94%) | 42 | (1.84%) | 4 | (0.18%) | NA |  | 435 | (19.05%) | 2,284 |
| NCI-H460 Nt.BspQI | 771 | (67.10%) | 65 | (5.66%) | 4 | (0.35%) | 5 | (0.44%) | 304 | (26.46%) | 1,149 |
| NCI-H460 DLE-1 | 1,738 | (78.08%) | 54 | (2.43%) | 3 | (0.13%) | 31 | (1.39%) | 400 | (17.97%) | 2,226 |
| PANC-1 | 651 | (58.65%) | 57 | (5.14%) | 3 | (0.27%) | 4 | (0.36%) | 395 | (35.59%) | 1,110 |
| SK-N-MC | 1,220 | (54.32%) | 316 | (14.07%) | 0 | (0%) | 8 | (0.36%) | 702 | (31.26%) | 2,246 |
| SK-BR-3 | 1,781 | (72.11%) | 40 | (1.62%) | 7 | (0.28%) | 75 | (3.04%) | 567 | (22.96%) | 2,470 |
| T47D Nt.BspQI | 1,188 | (67.85%) | 153 | (8.74%) | 16 | (0.91%) | 15 | (0.86%) | 379 | (21.64%) | 1,751 |
| T47D DLE-1 | 1,675 | (74.58%) | 44 | (1.96%) | 8 | (0.36%) | 43 | (1.91%) | 476 | (21.19%) | 2,246 |
| 261T | 1,636 | (87.21%) | 15 | (0.80%) | 2 | (0.11%) | NA |  | 223 | (11.89%) | 1,876 |
| 3096 | 1,758 | (83.40%) | 5 | (0.24%) | 0 | (0%) | NA |  | 345 | (16.37%) | 2,108 |
| 3717 | 1,809 | (81.52%) | 9 | (0.41%) | 0 | (0%) | NA |  | 401 | (18.07%) | 2,219 |
| 7052 Cancer | 1,768 | (80.04%) | 15 | (0.68%) | 0 | (0%) | NA |  | 426 | (19.28%) | 2,209 |
| 7052 Non-cancer | 1,704 | (88.43%) | 10 | (0.52%) | 0 | (0%) | NA |  | 213 | (11.05%) | 1,927 |
| 7403 Cancer | 1,752 | (82.88%) | 9 | (0.43%) | 0 | (0%) | NA |  | 353 | (16.70%) | 2,114 |
| 7403 Non-cancer | 1,757 | (88.38%) | 10 | (0.50%) | 0 | (0%) | NA |  | 221 | (11.12%) | 1,988 |
| 7518 | 1,759 | (82.93%) | 15 | (0.71%) | 0 | (0%) | NA |  | 347 | (16.36%) | 2,121 |
| 7528 Cancer | 1,812 | (84.28%) | 13 | (0.60%) | 0 | (0%) | NA |  | 325 | (15.12%) | 2,150 |
| 7528 Non-cancer | 1,810 | (88.34%) | 16 | (0.78%) | 0 | (0%) | NA |  | 223 | (10.88%) | 2,049 |
| 7622 Cancer | 1,773 | (82.01%) | 7 | (0.32%) | 0 | (0%) | NA |  | 382 | (17.67%) | 2,162 |
| 7622 Non-cancer | 1,729 | (84.42%) | 11 | (0.54%) | 0 | (0%) | NA |  | 308 | (15.04%) | 2,048 |
| 7708 | 1,736 | (84.64%) | 6 | (0.29%) | 0 | (0%) | NA |  | 309 | (15.07%) | 2,051 |
| 10974 | 1,780 | (75.39%) | 22 | (0.93%) | 1 | (0.04%) | NA |  | 558 | (23.63%) | 2,361 |
| 14369 | 1,748 | (76.30%) | 16 | (0.70%) | 1 | (0.04%) | NA |  | 526 | (22.96%) | 2,291 |
| Shuffle DLE-1 | 6,172 | (13.71%) | 3,640 | (8.08%) | NA |  | NA |  | 35,213 | (78.21%) | 45,025 |
| Shuffle Nt.BspQI | 2,131 | (22.17%) | 1,349 | (14.03%) | NA |  | NA |  | 6,134 | (63.80%) | 9,614 |

**Table S4:** Overlap of the SVs identified from the tongue squamous cell carcinoma samples and matched blood samples from the same patients. Each entry is the number of SVs commonly called from the two samples corresponding to the row and the column of the entry divided by the sample corresponding to the row, i.e. *x_ij_* = *|S_i_ ∩ S_j_|/|S_i_|*, where *x_ij_*is the entry in the *i*-th row and the *j*-th column of the table, *S_i_*and *S_j_*are the SV lists of the *i*-th and *j*-th samples, respectively, and *|S|* denotes the number of SVs in the set *S*. The largest value in each row is underlined.

|  | 7052 Cancer | 7052 Non-cancer | 7403 Cancer | 7403 Non-cancer | 7528 Cancer | 7528 Non-cancer | 7622 Cancer | 7622 Non-cancer |
| --- | --- | --- | --- | --- | --- | --- | --- | --- |
| 7052 Cancer | - | <u>84.01%</u> | 63.03% | 61.13% | 62.66% | 60.16% | 61.41% | 57.02% |
| 7052 Non-cancer | <u>86.76%</u> | - | 64.25% | 63.47% | 64.25% | 62.45% | 63.28% | 59.17% |
| 7403 Cancer | 63.17% | 62.75% | - | <u>85.29%</u> | 62.89% | 60.80% | 62.98% | 58.99% |
| 7403 Non-cancer | 64.84% | 64.89% | <u>89.25%</u> | - | 64.99% | 64.01% | 64.79% | 62.50% |
| 7528 Cancer | 62.82% | 62.91% | 62.96% | 61.64% | - | <u>83.46%</u> | 62.82% | 59.80% |
| 7528 Non-cancer | 66.19% | 66.80% | 65.94% | 66.24% | <u>90.03%</u> | - | 66.24% | 65.49% |
| 7622 Cancer | 59.45% | 59.41% | 60.58% | 58.91% | 60.09% | 58.27% | - | <u>77.23%</u> |
| 7622 Non-cancer | 63.12% | 63.80% | 65.52% | 65.36% | 65.67% | 66.40% | <u>88.59%</u> | - |

**Table S5:** List of parameters of COMSV and their default values. The order of the parameters follows their descriptions in the Materials and Methods section. *d_ref_* denotes the distance of the reference allele.

| Parameter | Description | Default value |
| --- | --- | --- |
| $a$ | Maximum percentage of observed-to-expected distance ratios strongly deviated from one for high-confidence alignments | 75 |
| $\gamma$ | Parameter for defining an observed-to-expected distance ratio to be strongly deviated from one | $\max(\delta, s_{min}/d_{ref})$ |
| $N$ | Minimum number of alignments to form a reinforcement group | 5 |
| $p_d$ | Minimum percentage of alignments with distance ratios strongly deviated from one for calling a suspected indel case | 10 |
| $\delta$ | Minimum ratio of distance change to support an indel | 0.05 |
| $s_{min}$ | Minimum size of an indel | 2,000bp |
| $n_s$ | Minimum size for a cluster to be retained | 3 |
| $c$ | Minimum number of aligned optical maps covering an indel region | 10 |

**Table S6:**
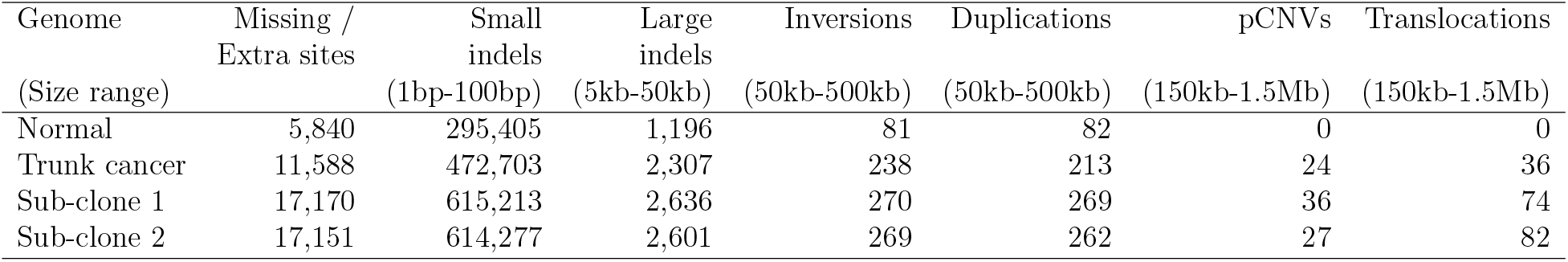
Numbers of small and large variations introduced to each genome of the second simulation experiment. Missing/Extra sites are damaged/novel labels caused by single nucleotide variations. pCNV: copy number variations due to large dissociative molecule fragments that cause polyploidy in cells.

**Table S7:** Sources of previously reported SVs identified from samples involved in this study using non-optical mapping data. Other data include PCR amplifications and the detection of SV-induced fusion genes based on RNA-seq. NA: not available

| Sample ID | Short Reads | 10x linked-reads | Hi-C | Other data |
| --- | --- | --- | --- | --- |
| A549 | [10] | NA | [10] | NA |
| Caki2 | [10] | NA | [10] | [10] |
| Huh7 | NA | NA | NA | [37] |
| K562 | [10] | NA | [10] | [10] |
| LNCaP | [10] | NA | [10] | NA |
| NCI-H460 | [10] | NA | [10] | NA |
| PANC-1 | [10] | NA | [10] | NA |
| SK-N-MC | [10] | NA | [10] | NA |
| SK-BR-3 | [38] | [38] | NA | [38] |
| T47D | [10] | NA | [10] | [10] |

### S2 Supplementary figures

**Figure S1:**
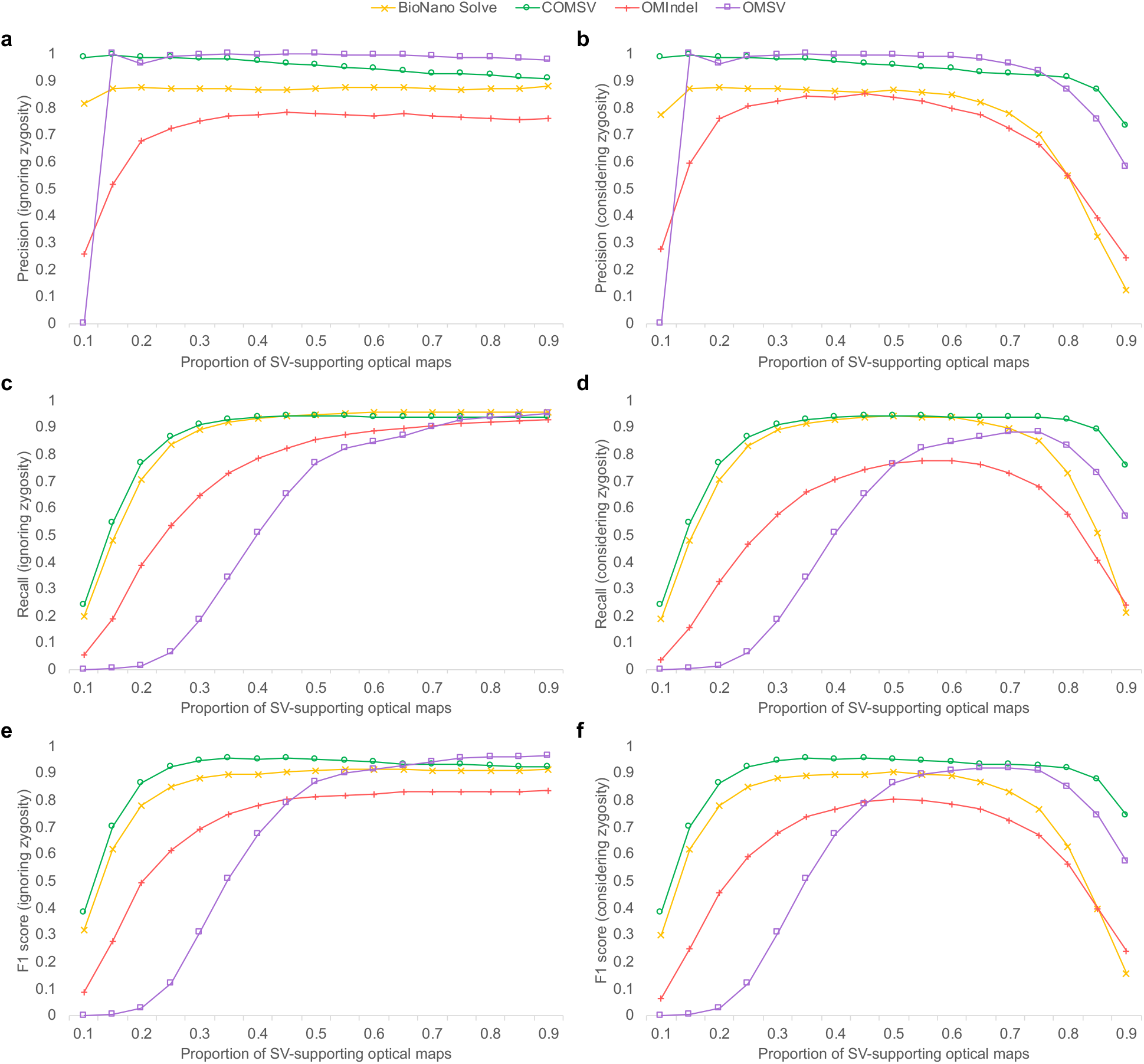
Comparisons between the performance of COMSV and three existing SV calling methods based on simulated data with a varying proportion of supporting optical maps at the SV loci. Precision (**a, b**), recall (**c, d**) and F1 score (**e, f**) of the different methods when the zygosity of the SVs are ignored (a, c, e) or considered (b, d, f) by the performance measures.

**Figure S2:**
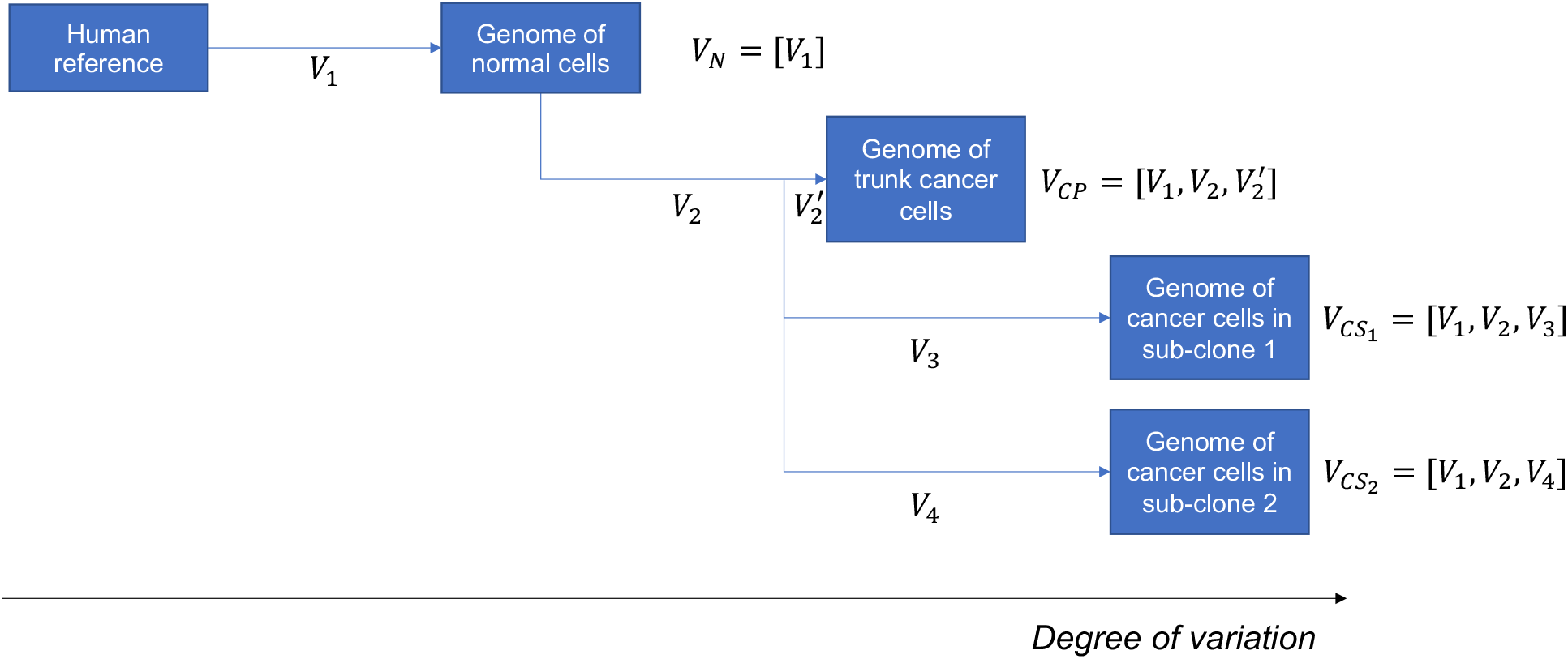
The evolution of cells in the second simulation experiment. Based on the human reference, a first set of SVs (*V*_1_) were generated to create the genome of the normal cells. A second set of SVs (*V*_2_) were then generated to create the genome of trunk cancer cells. While the trunk cancer cells continued to evolve with a small additional set of SVs (*V* ^t^), two cancer sub-clones were evolved with their own sets of SVs (*V*_3_ and *V*_4_, respectively). The final sets of SVs contained in the genomes of the normal cells, trunk cancer cells, cells in the first sub-clone and cells in the second sub-clone are denoted as *V_N_*, *V_CP_*, *V_CS_*_1_ and *V_CS_*_2_, respectively.

**Figure S3:**
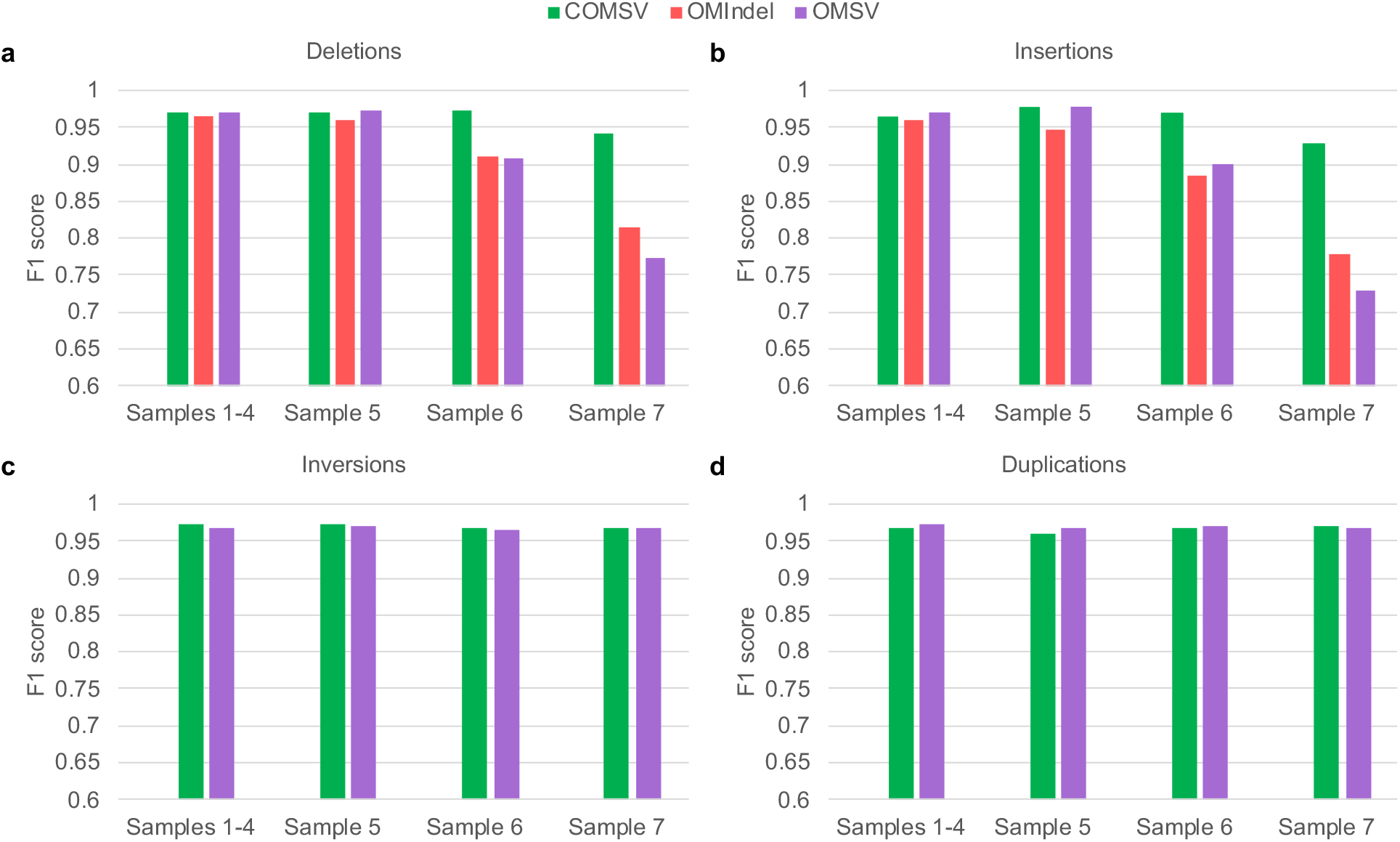
Comparisons between the performance of COMSV and other existing SV calling methods based on simulated data with different cell compositions, with the correct optical map alignments supplied as inputs. **a-d** For each type of SVs, including deletions (a), insertions (b), inversions (c) and duplications (d), the performance of each method when applied to the homogeneous cell samples with one cell type per sample (Samples 1-4) and the heterogeneous cell samples with multiple cell types per sample (Samples 5-7) is shown. Since Bionano Solve could not be run with optical map alignments supplied as inputs, it was omitted from these comparisons.

**Figure S4:**
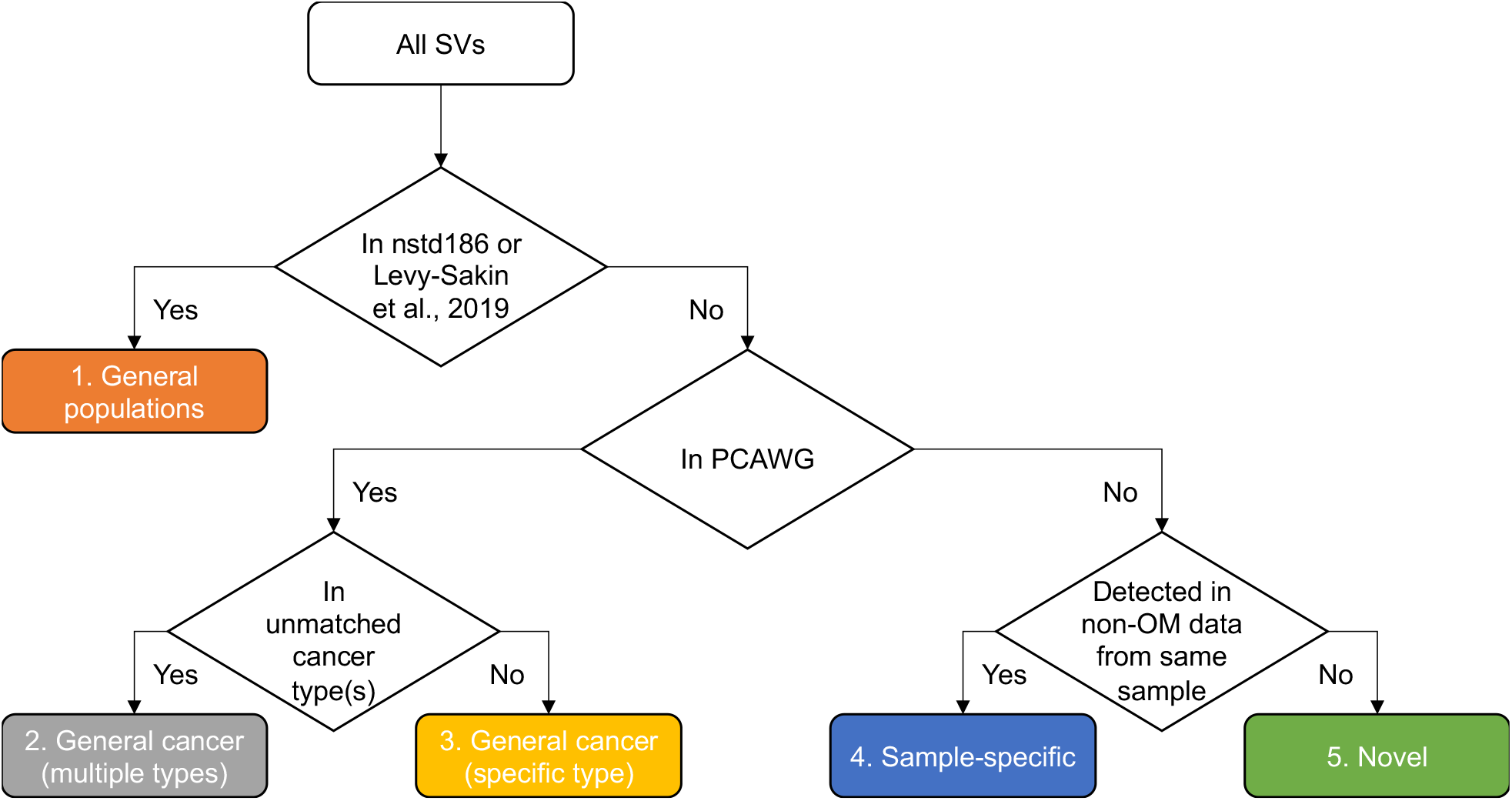
Classification of SVs identified from the real samples based on supports provided by independent data.

**Figure S5:**
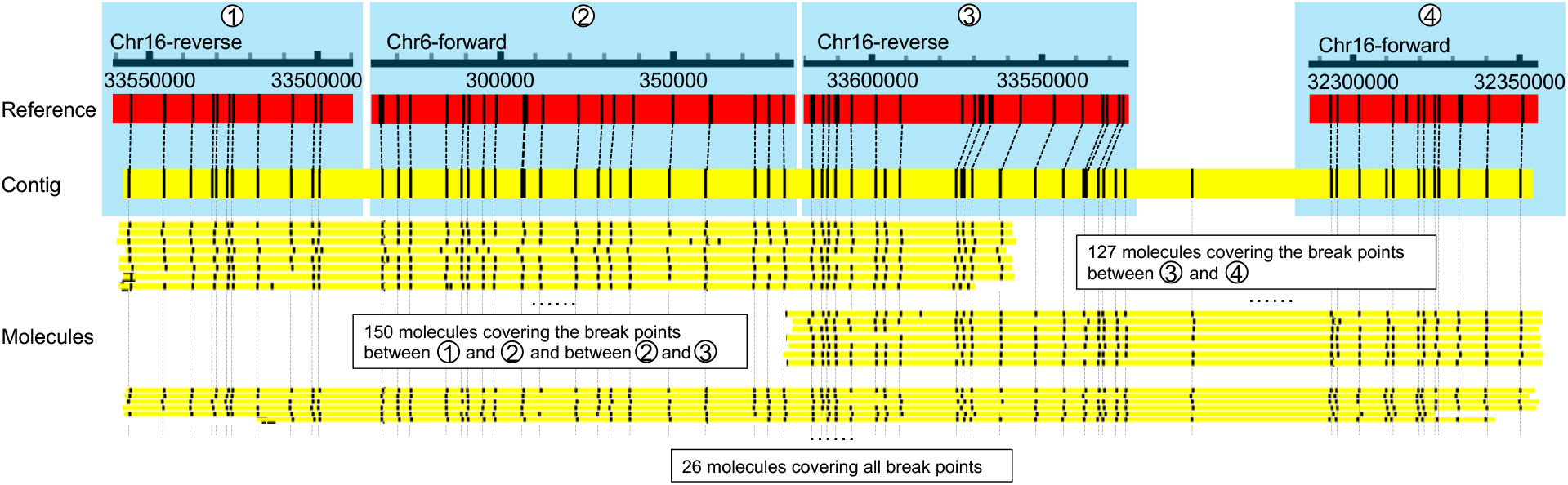
Supporting evidence for a complex SV-containing contig. The alignment between the reference (red bars), optical map contig (thick yellow bar) and individual optical map molecules (thin yellow bars) are shown for the region highlighted in a box with dashedline boundaries in Figure 6d. Aligned segments (1-4) are numbered in the same way as in Figure 6d.

**Figure S6:**
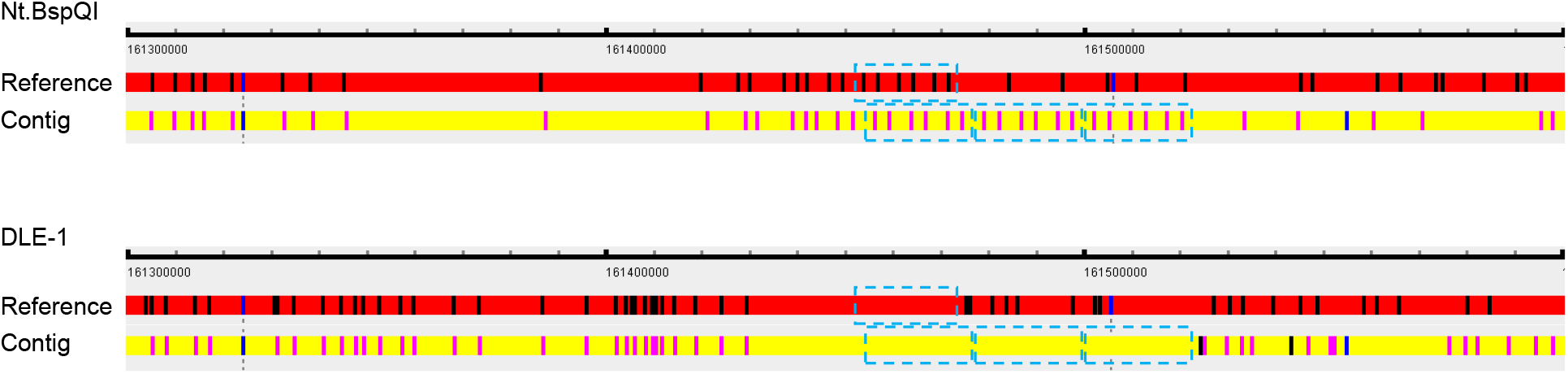
A duplication identified from Nt.BspQI data but not DLE-1 data. The duplication, identified from LNCaP, displays a pattern (highlighted by the blue boxes) that repeats three times in Nt.BspQI data. DLE-1 labels are completely missing in this region.

**Figure S7:**
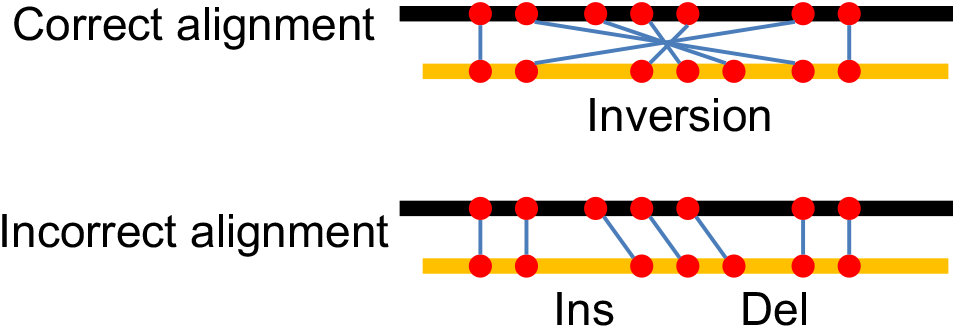
Illustration of a situation that leads to missed call of inversions. The upper part show the correct alignment of the labels on a molecule (yellow bar) with those on a reference (black bar), which should support an inversion. However, due to the symmetric pattern of the labels in the inverted region, the actual alignment produced can be the one shown in the lower part, which instead supports an insertion and a deletion.

### S3 Supplementary files

Supplementary File 1: List of SVs identified from individual human samples

Supplementary File 2: Unified list of non-redundant novel SVs identified from the different human samples

Supplementary File 3: Lists of genomic regions excluded from SV calling

